# Mutation in shoot-to-root mobile transcription factor, ELONGATED HYPOCOTYL 5, leads to low nicotine levels in tobacco

**DOI:** 10.1101/2022.01.05.475064

**Authors:** Deeksha Singh, Shambhavi Dwivedi, Hiteshwari Sinha, Nivedita Singh, Prabodh Kumar Trivedi

## Abstract

Tobacco remains one of the most commercially important crops due to the parasympathomimetic alkaloid nicotine used in cigarettes. Most genes involved in nicotine biosynthesis are expressed in root tissues; however, their light-dependent regulation has not been studied. We identified the ELONGATED HYPOCOTYL 5 homolog, NtHY5, from *Nicotiana tabacum* and demonstrated its role in nicotine biosynthesis. We report the development of CRISPR/Cas9-based mutant plants, *NtHY5^CR^*, and show down-regulation of the nicotine biosynthetic pathway, whereas NtHY5 overexpression (NtHY5OX) plants show the opposite effect. Grafting experiments using wild type, *NtHY5^CR^,* and NtHY5OX indicated that NtHY5 moves from shoot-to-root to regulate nicotine biosynthesis in the root tissue. We conclude that shoot HY5, directly or through enhancing expression of the HY5 in the root, promotes nicotine biosynthesis. The CRISPR/Cas9-based mutants developed, in this study; with low nicotine accumulation in leaves could help people to overcome their nicotine addiction and the risk of death from tobacco use.

## INTRODUCTION

Nicotine is the most abundant alkaloid in *Nicotiana tabacum* (Dewey and Xie, 2013), contributing to 90% of the total alkaloids. The development of tobacco plants with significantly less nicotine using genetic engineering techniques is urgently needed to reduce the death rate and diseases caused by tobacco consumption, as the tobacco epidemic is one of the biggest threats to public health, killing millions of people every year around the world (WHO, 2015, 2021). Nicotine is structurally composed of pyridine and pyrrolidine rings (Hibi et al., 1994). The biosynthesis of nicotine occurs in the cortical cells of roots through various enzymatic steps (**Figure 1A**) and is cloistered in vacuoles (Shoji et al., 2009). It is further transported to leaves via xylem (Dewey et al., 2013; Shoji et al., 2021; Hayashi et al., 2020), where it is used for defence against herbivory by the plant (Steppuhn et al., 2004). Quinolinate phosphoribosyl transferase (QPT) (Sinclaie et al., 2000) acts on nicotinic acid and catalyze the formation of the pyridine ring (Hashimotoet et al., 1994). The pyrrolidine ring, on the other hand, is formed from putrescine derived directly from ornithine by ornithine decarboxylase (ODC) (Imanshi et al., 1998). Putrescine N-methyltransferase (PMT) converts putrescine to N-methyl-putrescine (Riechers et al., 1999), which is further oxidized by N-methyl-putrescine oxidase (MPO) and cyclized to N-methyl pyrrolinium cation (Naconsie et al., 2014). The cation finally condenses with nicotinic acid catalysed by NADPH-dependent PIP reductase A622 (DeBoer et al., 2009) and the berberine bridging enzyme-like (BBL) proteins to form nicotine (Kajikawa et al., 2011; Dewey et al., 2013; Hibi et al., 1994; Shoji et al., 2021; Shoji et al., 2011).

**Figure 1.**
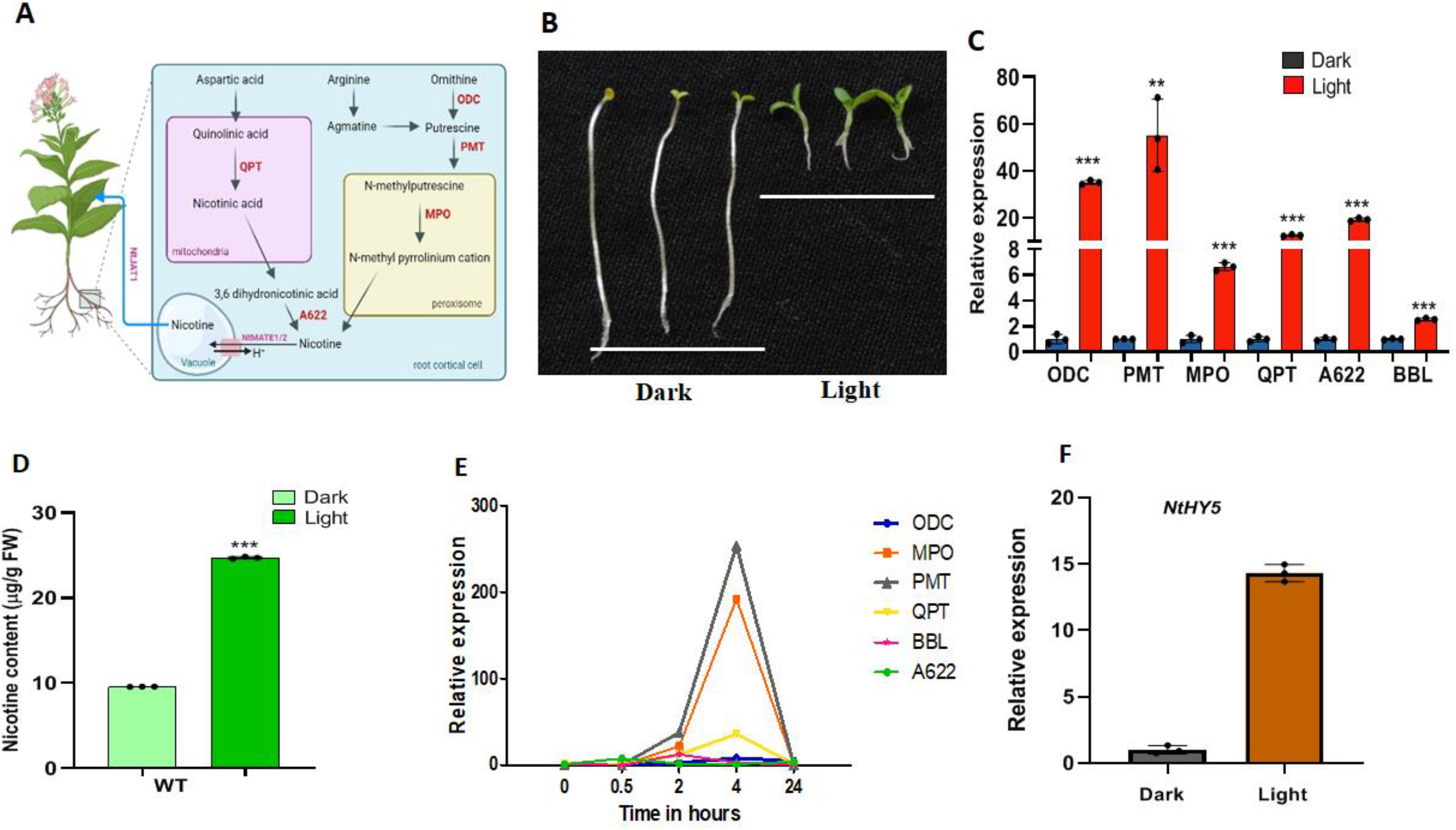
Light-mediated modulation of nicotine and flavonol content in WT tobacco. A, Diagrammatic representation of nicotine pathway of tobacco. B, Phenotype of 10-d-dark and light grown WT seedlings. C, Expression analysis of nicotine pathway genes (*ODC, PMT, MPO, QPT and A622*) in WT seedlings grown in light (root were covered to inhibit light illumination) and dark for 10-days through qRT-PCR. D, Quantitative estimation of nicotine in 10-day-dark and light grown seedlings (using only hypocotyl region) through HPLC. E, Time dependent qRT-PCR expression analysis of nicotine pathway genes (ODC, MPO, PMT, QPT and A622) in 10-day dark grown WT tobacco seedlings subsequently transferred to light for 0.5, 2, 4 and 24 hours. F, Relative expression of NtHY5 in 10-d-dark and light grown seedlings. Statistical analysis was performed using two-tailed Student’s t-test. Error bars represent SE of means (n=3) For hypocotyl and root length (n=15). Tubulin was used as endogenous control to normalize the relative expression levels. Error bars represent standard deviation. Asterisks indicate a significant difference, *P < 0.1, **P < 0.01, ***P < 0.001.

Previous research has suggested that a number of regulatory factors are involved in the production of nicotine. According to one study, there are two AP2/ERF genes, NtORC1/ERF221 and NtJAP1/ERF10, both of them positively control the PMT gene. In contrast, NtORC1/ERF221 overexpression only causes high nicotine alkaloid levels (De Boer et al., 20011). Microarray analysis using *Nicotiana tabacum* NIC2 mutant identified several AP2/ERF genes which can regulate nicotine biosynthesis. Seven clustered AP2/ERF genes at the NIC2 locus belong to the group IX AP2/ERF subfamily. One of them, NtERF189, encodes for subclade 2-1 of group IX AP2/ERF protein, is MeJA sensitive, and was expressed in roots, where nicotine is primarily synthesized. The master regulatory role of NtERF189 in nicotine biosynthesis in tobacco is suggested by the fact that knockout mutant of NtERF189 and NtERF199 synthesize reduced nicotine (Hayashi et al., 2020). The NtERF189 protein recognizes GCC-box-like elements, (A/C)GC(A/C)(A/C)NCC, in the promoter of nicotine biosynthetic enzyme genes (Shoji et al., 2012). NtERF32 (EREBP2), another AP2/ERF gene, has also been discovered, which is not a part of the NIC2 locus. Analysis suggested that NtERF32 may act as a transcriptional activator to regulate the expression of multiple genes involved in nicotine synthesis by interacting with the GCC-box-like element (Sears et al., 2014). According to one study, MYC2-type bHLH transcription factors play a role in regulating the biosynthesis of nicotine. While their overexpression marginally boosted PMT expression and nicotine content, RNA silencing of NbbHLH1 and NbbHL2 significantly lowered the expression levels of the genes encoding the nicotine biosynthesis enzymes. The target GCC-box of NtORC1/ERF221, where NbnHLH1 can bind, is located proximal to the G-box in the PMT promoter. By interactions with both components, NtORC1/ERF221 and NbbHLH1 work together to control PMT expression. Moreover, *N. tabacum* has been found to contain homologous genes of NbbHLH1 and NbbHLH2. The transcript levels of NIC2-locus AP2/ERF genes, transporter genes, and nicotine biosynthesis enzymes are favorably regulated by NtMYC2a and NtMYC2b (Shoji et al., 2011). The direct interaction between NtMYC2 and NtJAZ1 has also been confirmed. The basic helix-loop-helix protein MYC2 and an orthologous pair of ethylene response factor (ERF) family proteins, ERF189 and ERF199, form a cascade to regulate the nicotine production pathway by specifically recognizing the GC-rich sequences in the promoter regions of their downstream genes (Shoji et al., 2010; Shoji et al., 2013; Shoji and Yuan et al., 2021). NtMYB305a has recently been revealed to bind to the JA-responsive GAG region in the NtPMT1a promoter to positively regulate nicotine biosynthesis (Bian et al., 2022). Despite the fact that these studies have been done on transcriptional regulation of nicotine biosynthesis, the light-dependent regulation of the nicotine pathway in tobacco is not explored yet, although studies suggested the presence of two functionally important *cis*-elements, a G-box element and a GCC-box element, (Shoji et al., 2000; Oki and Hashimoto et al., 2004; Xu and Timko, 2004) in the promoter region of PMT.

Light plays a crucial role in plant developmental processes such as flowering, seed development, organ development, seed germination and seedling de-etiolation (Chory et al., 2010; Vanhaelewyn et al., 2016; Gangappa et al., 2016; Bhatnagar et al., 2020). Many transcription factors of various families, including bHLH, bZIP, MYB, GATA and Zinc-finger are activated after light perception by photoreceptors (Chory et al., 2010). Constitutive nuclear-localized ELONGATED HYPOCOTYL 5 (HY5), a bZIP family transcription factor, acts downstream to the photoreceptors and plays a central role in seedling development via integrating light and phytohormone signaling pathways (Koornneef et al., 1980; Osterlund et al., 2000; Ulm et al., 2012; Li et al., 2013). In *Arabidopsis*, HY5 acts as a master regulator by binding to *cis*-regulatory elements of more than 3000 genes directly (Zhang et al., 2011) and regulates flavonoid and terpenoid biosynthesis (Bhatia et al., 2018; Michael et al., 2021). Though extensive characterization of HY5 and flavonoid biosynthesis has been done in several plant species, but, there are no reports of characterization of *N. tabacum* HY5 and its role in light-regulated photomorphogenesis as well as nicotine and flavonoid biosynthesis in the plant.

In this study, to decipher the function of HY5 of tobacco in nicotine and flavonoid biosynthesis, we identified and functionally characterized NtHY5 through complementing *Arabidopsis* HY5 mutant (*athy5*), developing NtHY5-overexpressing lines and CRISPR/Cas9-based mutants. In our study, through different combinations of grafting experiments, we showed that in response to light, NtHY5 moves from shoot-to-root and may activate the expression of *NtHY5* in roots as well as bind directly to the promoter of genes involved in nicotine biosynthesis. CRISPR/Cas9-based HY5 mutants accumulated significantly lower amounts of nicotine compared to wild-type tobacco plants, suggesting the involvement of light-associated signaling in the regulation of nicotine biosynthesis in roots.

## RESULTS

### Identification of light-associated signaling component involved in nicotine biosynthesis

Based on the presence of G and GCC box in the *NtPMT* promoter described in the previous studies (Xu et al., 2017; Shoji et al., 2000; Oki and Hashimoto et al., 2004; Xu and Timko et al., 2004), we hypothesized that light could be the factor affecting the nicotine biosynthesis in tobacco. Previous studies have also established that HY5 binds to G-box (CACGTG) in the promoters of light-responsive genes (Zhang et al., 2011). To validate the hypothesis, tobacco seedlings were grown in the light, keeping roots in the dark to maintain natural conditions, and continuously dark for 10 days. Phenotypically, light- and dark-grown seedlings showed substantial differences in hypocotyl length, cotyledon opening and chlorophyll content (**Figure 1B**). The gene expression analysis in roots for nicotine pathway genes, and seedlings for flavonoid pathway genes, suggested a higher expression of genes involved in the nicotine biosynthesis pathway (*NtODC, NtPMT, NtMPO, NtQPT and NtA622*) (**Figure 1C**) and flavonoid biosynthesis (*CHS, CHI, FLS and DFR*) (**Supplemental Figure S1A**) in light-grown compared to dark-grown seedlings. Expression of the regulatory gene of the flavonoid pathway, *NtMYB12*, was also higher in light-grown than dark-grown seedlings (**Supplemental Figure S1A**). We measured the nicotine content in the hypocotyl region of seedlings, and analysis suggested significantly higher nicotine content in light-grown compared to dark-grown seedlings (**Figure 1D**). We also evaluated the content of flavonols (kaempferol, quercitin, and rutin), another significant secondary metabolite in tobacco. Analysis suggested a significantly higher accumulation of flavonols (**Supplemental Figure S1B**) in light-grown seedlings compared to dark-grown seedlings. Additionally, 10-day-dark grown seedlings were exposed to light for 0.5, 2, 4, and 24 hours to observe the light-responsiveness of nicotine pathway genes. The analysis suggested enhanced expression of most of the genes until 4 hours of light exposure. *NtMPO* and *NtODC* showed the maximum induction (**Figure 1E**).

As HY5 is a key light signaling component in several plants (Nawkar et al., 2017; Burman et al., 2018), we identified the tobacco HY5, NtHY5, and investigated its function in nicotine and flavonol accumulation. NtHY5 showed more than 90% sequence similarity with *Arabidopsis* HY5, AtHY5. Sequence analysis of NtHY5 suggested that it contains 4 exons and 3 introns similar to *AtHY5* (**Supplemental Figure S2A**). Amino acid sequence analysis suggested that NtHY5 contains leucine-rich zinc finger motif and the VPE motif (COP1-interaction motif) (Holm et al., 2002) and has a high degree of similarity with SlHY5 and AtHY5 (**Supplemental Figure S2B**). The conserved domain database analysis revealed that NtHY5 is a member of the bZIP superfamily, similar to HY5 proteins in other plant species (**Supplemental Figure S2C**). Phylogenetic and pairwise distance analysis suggested that NtHY5 is closer to CaHY5 and SlHY5 than AtHY5 (**Supplemental Figure S3**).

The changes in phenotype, expression of genes and metabolite content in the dark- and light-grown seedlings led us to quantify the expression of identified NtHY5 gene in the dark- and light-grown seedlings. The expression of NtHY5 was significantly low in dark-grown seedlings compared to light-grown seedlings (**Figure 1F**). As the content of flavonols changes in response to light (Pandey et al., 2014; Bhatia et al., 2018), we studied the responsiveness of flavonols (kaempferol and quercitin) towards light after exposing 5-day-old dark-grown WT seedlings to light conditions for 3 h. The analysis suggested a higher accumulation in dark-to-light exposed seedlings compared to only dark-grown seedlings (**Supplemental Figure S4**).

To investigate the spatio-temporal expression of *NtHY5* and nicotine pathway genes, a comparative analysis was carried out at different stages of tobacco development. The analysis suggested 4- to 5-fold higher expression of NtHY5 (**Supplemental Figure S5A**) in the leaf, compared with the seedling, root, stem, and flower of the plant. Expression of *NtODC, NtQPT, NtPMT, NtMPO* and *NtA622* was several folds higher in root in comparison to leaf, flower, seedling, and stem (**Supplemental Figure S5B**).

### NtHY5 interacts with the promoters of genes involved in nicotine and flavonoid biosynthesis

According to the aforementioned data, structural genes of the nicotine biosynthesis (*NtPMT*, *NtQPT*, and *NtODC*) and flavonol regulatory genes (*NtMYB12*) have substantially higher expression in the light compared to dark. It was, therefore, intriguing to analyze whether or not the light signaling component, NtHY5, directly regulates these genes. *in-silico* analysis suggested that promoters of *NtPMT*, *NtQPT*, *NtODC*, and *NtMYB12* contain potential LREs (GATA, BOX-4, ACE, and LTR) upstream of the transcription start site (**Supplemental Table S1-S4**). Analysis suggested the presence of G-boxes with core motif ACGT at -168 bp (G-box I) and -1040 bp (G-box II) in the *NtPMT* promoter, -178 bp (G-box I) upstream of the *NtODC* transcription start site and -36 bp (G-box I) upstream of the *NtQPT* promoter from the transcription start site (**Figure 2A**). NtMYB12 contains G-boxes at -1249 bp and -207 bp upstream of the transcription start site (**Supplemental Figure S6A**).

**Figure 2.**
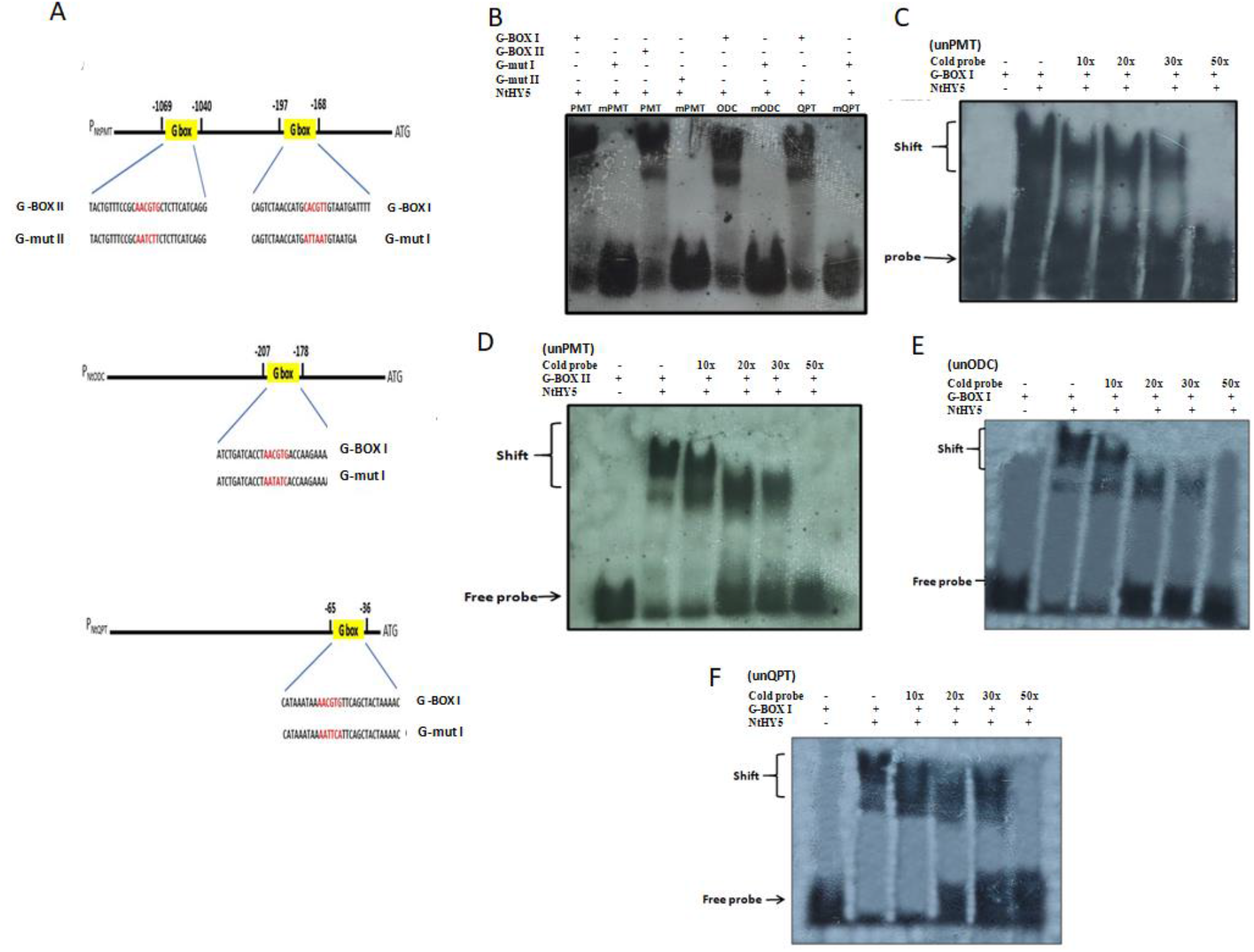
In vitro interaction between NtHY5 and the ACGT motif of G-BOX present at the promoters of *NtPMT, NtODC and NtQPT* genes. A, Representation of the probes (G-BOXI, G-mut I, G-BOXII and G-mut II) designed upstream of the transcription start site of NtPMT, NtODC and NtQPT. G-BOX is altered by multiple base substitutions. B, EMSA (Electrophoretic mobility shift assay) for the binding of 6X-His-HY5 (NtHY5) with core ACE-DIG element (HY5 binding site that is G-BOXI and G-BOXII) present in the LRE motifs of the NtPMT, NtODC and NtQPT respectively. Upper and lower arrows indicate shift and free probe respectively. C-F, EMSA with competition between digoxigen in labelled G-BOXI, unlabeled G-BOX I (cold), G-BOXII and unlabeled G-BOX II (cold) probe to bind to 6X-His-HY5 respectively. Superscript values 10X, 20X, 30X and 50X represent increasing amount of cold probe NtPMT-P1, NtPMT-P2, NtQPT and NtODC respectively. Upper and lower arrows indicate shift and free probe, respectively.

An electrophoretic mobility shift assay (EMSA) was employed to analyze the *in vitro* interaction between ACGT (core motif of G-boxes) and recombinant 6XHis-tagged NtHY5 (NtHY5His) expressed in *E. coli*. In the EMSA with DIG-labeled G-box probe (G-BOX I and G-BOX II for PMT and G-BOX I for ODC and QPT) (**Figure 2B**) and NtMYB12 probe (G-BOX I and G-BOX II) (**Supplemental Figure S6B**), a band shift was observed upon addition of the NtHY5-His protein. At the same time, no band shift was observed with the mutated probe (mPMT, mODC and mQPT). In the competition assay, as the concentration of unlabeled probe or cold probe (unPMT, unODC, unQPT) (**Figure 2C-F**) and (unNtMYB12) (**Supplemental Figure S6C and D**) increases from 10X to 50X, the intensity of binding got diminished. These findings suggested that NtHY5 interacts with promoters of *NtPMT*, *NtODC*, and *NtQPT* to regulate the expression of these genes leading to the regulation of nicotine content, whereas to regulate flavonoid content, it binds to NtMYB12.

### NtHY5 is a functional ortholog of AtHY5

To investigate the biological function of NtHY5, the homozygous transgenic lines in the *Arabidopsis* HY5 mutant (*athy5* background) were generated and used for detailed analysis. Phenotypic complementation by NtHY5 in terms of hypocotyls length, root length and rosette diameter was analysed (**Figure 3A**). The average hypocotyl lengths of 7-d-old wild-type (WT), *hy5* mutant, and NtHY5;*hy5* seedlings were approximately 0.2, 0.7, and 0.4 mm, respectively, demonstrating that complemented lines have recovered their lost hypocotyl length (**Figure 3B**). As AtHY5 plays a significant role in light-dependent root growth in *Arabidopsis* (Zhang et al., 2017), the root length of 7-day-old light-grown WT, *hy5* mutant, and NtHY5;*hy5* complemented *Arabidopsis* seedlings were measured. Analysis suggested that the root length of complemented seedlings is recovered to WT seedlings (**Figure 3C**). The average rosette diameter of the *hy5* mutant was about 6 cm, whereas the diameters of the WT and complemented lines were 4 and 3 cm, respectively, demonstrating the recovery of rosette size as well (**Figure 3D, Supplemental Figure S7A**). Similarly, seeds of the *hy5* mutant also regained the size similar to WT after complementation (**Figure 3E and 3F**). These results suggest that NtHY5 could complement the phenotypic function of the AtHY5 in the *hy5* mutant.

**Figure 3.**
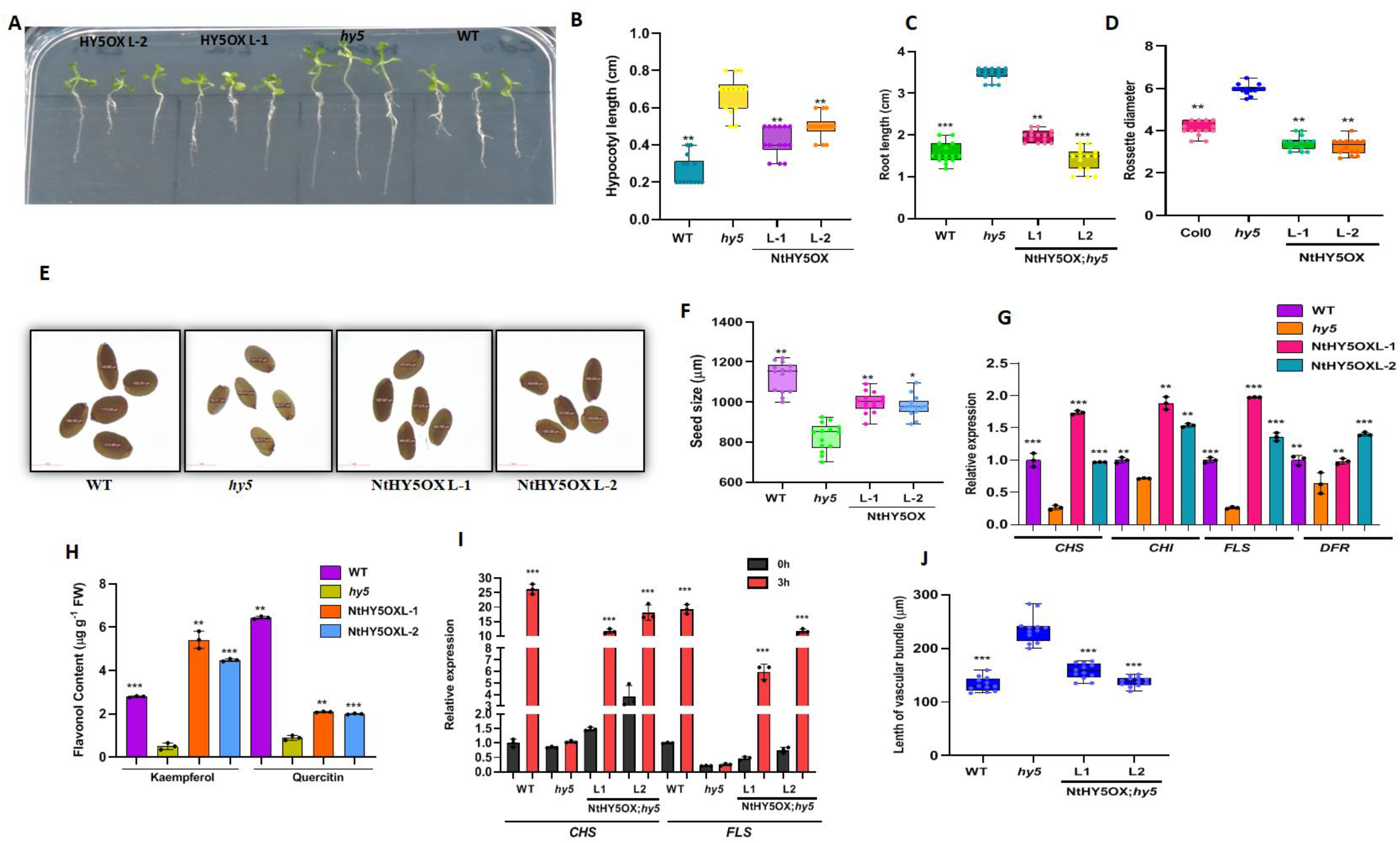
Complementation of *Arabidopsis hy5* mutant by NtHY5. **A**, Phenotype of 7-d-old light-grown WT (wild type; Col-0), *hy5* and NtHY5OX; *hy5* seedlings (L1 and L2 are two independent transgenic lines overexpressing NtHY5 in *hy5* mutant background). B, Hypocotyl length of 7-d-old white light grown WT, *hy5*, L1 and L2 seedlings. C, Root length of 7-d-old white light grown WT, hy5, L1 and L2. D, Rosette diameter of 30-day-old WT, *hy5*, L1 and L2 plants. E, Representative photographs of the seed size of WT, *hy5* and transgenic lines (L1 and L2). Bars = 1 mm. F, Measurement of seed size of 30-day-old WT, *hy5*, L1 and L2 plants. G, Relative expression of pheylpropanoid pathway genes (*AtCHS, AtCHI, AtFLS and AtDFR*) in 7-d-light grown WT, *hy5*, L1 and L2 seedlings. H, Quantitative estimation of flavonol (Kaempferol and Quercitin) content in 7-d-old light grown WT, *hy5*, L1 and L2 seedlings through HPLC. I, Expression analysis of *AtCHS, AtCHI, AtFLS* and *AtDFR* in WT, *hy5*, L1 and L2 seedlings grown for 5 days in dark followed by an exposure to light for 3hr. J, Length of vascular bundle of 30-day-oldstem of WT, *hy5*, L1 and L2 plants. Statistical analysis was performed using two-tailed Student’s t-test. Error bars represent SE of means (n=3). For hypocotyl and root length (n=10-12). Tubulin was used as endogenous control to normalize the relative expression levels. Error bars represent standard deviation. Asterisks indicate a significant difference, *P < 0.1, **P < 0.01, ***P < 0.001.

### NtHY5 can compensate for the loss at transcript and metabolite levels in *Arabidopsis hy5* **mutant**

To understand the function of NtHY5 at transcript and metabolite levels, the expression of flavonol biosynthetic pathway genes was studied in WT, *hy5* and NtHY5;*hy5* complemented seedlings. The analysis suggested that the expression of the NtHY5 in the *hy5* mutant background restored the expression of genes associated with the phenylpropanoid pathway (*AtCHS*, *AtCHI*, *AtFLS* and *AtDFR*) (**Figure 3G**). The levels of flavonols (kaempferol and quercetin) were also restored to WT in NtHY5;*hy5* lines compared with the *hy5* mutant **(Figure 3H)**. To determine the responsiveness of complemented lines towards light, expression of genes associated with flavonoid biosynthesis was study in 5-day-old dark-grown seedlings subsequently exposed to light for 3h. The transcript level of structural genes increased on exposure to light for 3h in complemented lines. By contrast, *hy5* mutant showed inhibited light-dependent induction of gene expression **(Figure 3I)**. In addition, expression of MYB family transcription factors also enhanced in NtHY5 complemented lines after 3h of light induction compared with *hy5* mutant seedlings **(Supplemental Figure S7B)**.

The phenylpropanoid pathway synthesizes various secondary metabolites, including flavonols, flavones, flavonols, isoflavones, flavan-3-ol, proanthocyanidins (PAs), anthocyanin and lignin. Phenylalanine is reported to be a common precursor for all of them (Vogt et al., 2010). Thus, we studied the effect of NtHY5 on lignin and anthocyanin production in *Arabidopsis* complemented lines. For lignin staining, the stem cross-sections of the WT, *hy5*, and NtHY5;*hy5* were used. The results showed a decreased lignification in the interfascicular and vascular tissues (**Supplemental Figure S7C**). Additionally, compared to WT and *hy5* mutant plants, NtHY5;*hy5* had a shorter vascular bundle in length (**Figure 3J**) this suggested the vascular damage due to lesser lignin. As AtHY5 is known to be involved in anthocyanin and chlorophyll biosynthesis (Oyama et al., 1997; Chattopadhyay et al., 1998), we estimated anthocyanin and chlorophyll contents of WT, *hy5*, and NtHY5;*hy5* seedlings. In 7-day-old light-grown seedlings, the total chlorophyll content of NtHY5;*hy5* was higher than WT and *hy5* (**Supplemental Figure S7D**). The anthocyanin content in NtHY5;*hy5* was higher than *hy5* seedlings but found to be comparatively lesser than WT (**Supplemental Figure S7E**). These findings imply that NtHY5 can compensate for the loss in the *athy5* mutant at the transcript and metabolite levels.

### Nicotine and flavonoid production in tobacco is regulated by NtHY5

To gain a deeper insight into NtHY5 function and its involvement in nicotine and flavonoid, we developed NtHY5 mutant plants using the CRISPR–Cas9 approach and NtHY5 overexpression (NtHY5OX) plants. To mutate NtHY5, we used guide RNAs (gRNAs) targeting the coding region of NtHY5 (**Supplemental Figure S8A**). The nucleotide sequence analysis of the mutants revealed deletions and insertions in the coding regions of NtHY5 (**Supplemental Figure S8B**). The mutants (*NtHY5^CR^* plants) showed frame-shift mutations leading to truncated peptides (**Supplemental Figure S8C**). In 20-day-old light-grown seedlings where roots were covered to inhibit light illumination to maintain natural condition (**Figure 4A**), qRT-PCR expression analysis revealed the downregulation of NtHY5 in mutants, while overexpression seedlings showed a higher expression (**Figure 4B**). We also analyzed the expression of nicotine pathway genes (*NtODC, NtPMT, NtMPO, NtQPT* and *NtA622*). The analysis suggested a significant downregulation of genes involved in nicotine biosynthesis in *NtHY5^CR^* seedlings, whereas overexpression seedlings showed an increase in expression in comparison to WT (**Figure 4C**). Additionally, structural genes of the phenylpropanoid pathway, such as *NtCHS, NtCHI, NtFLS*, and *NtDFR,* and regulatory gene, *NtNtMYB12,* were downregulated in *NtHY5^CR^* seedlings and the opposite was observed for overexpression lines (**Supplemental Figure S9A and B**).

**Figure 4.**
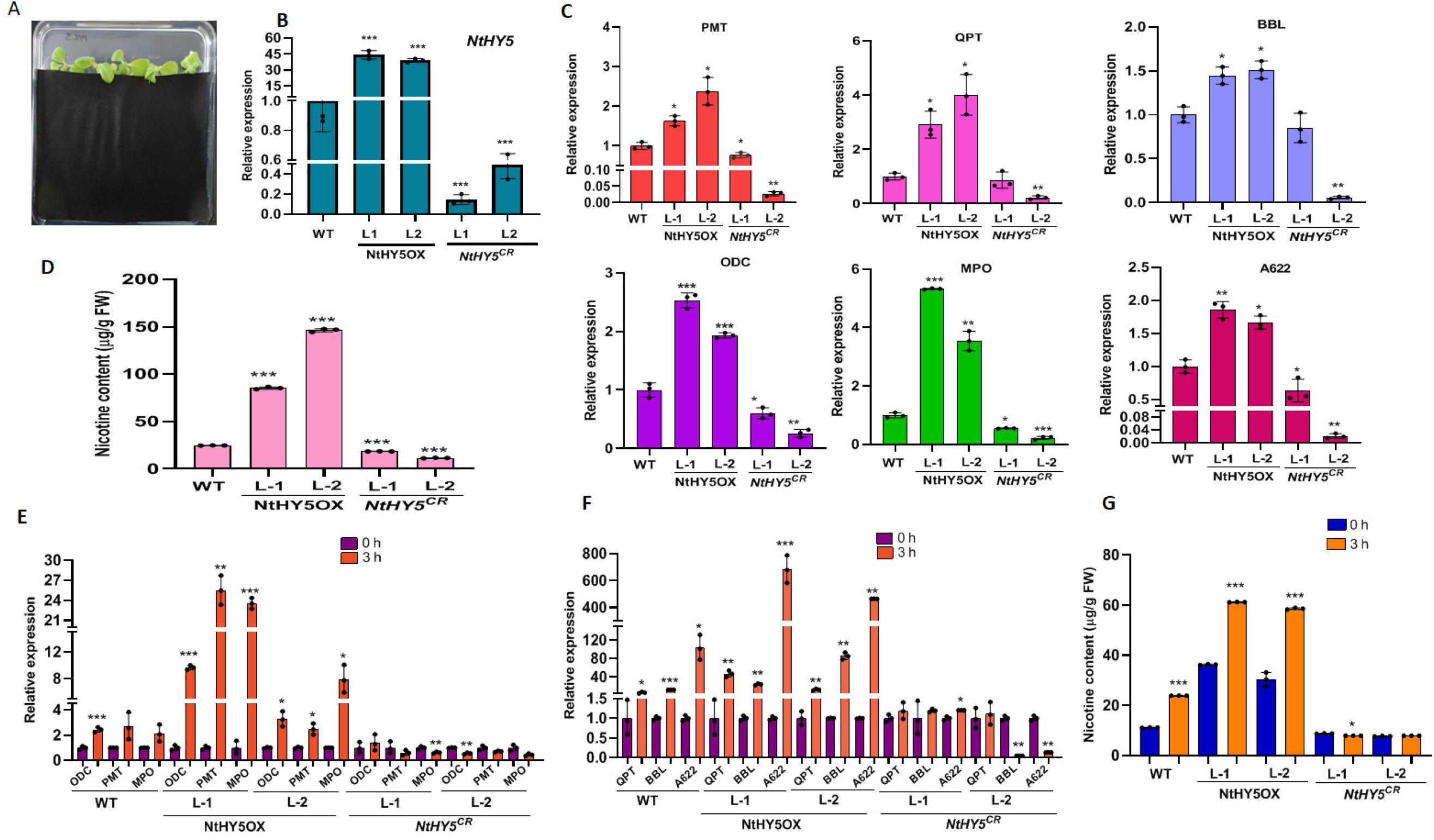
Overexpression of NtHY5 and CRISPR-Cas9-mediated knockout of NtHY5 modulates gene expression and accumulation of metabolite in tobacco seedlings. A, Visual representation of the 20-day-old seedlings with root covered with black paper after germination for 5 days to inhibit light exposure. Roots were covered with black paper for 14 days. B, Relative expression of NtHY5 in 20-day-old light grown WT, NtHY5OX and *NtHY5^CR^* lines of tobacco. B, C, Relative transcript abundance of nicotine pathway genes (*NtPMT, NtODC, NtQPT, NtMPO, NtBBL and NtA622*) in 20-day-old light grown WT, NtHY5OX and *NtHY5^CR^* lines. D, Quantification of nicotine content in 20-day light-grown WT, NtHY5OX and *NtHY5^CR^* leaves through HPLC. E and F, Relative transcript abundance of nicotine pathway genes (*NtODC, NtPMT, NtMPO, NtQPT, NtBBL* and NtA622) in 10-day-old dark-grown seedlings and seedlings transferred light for 3h in NtHY5OX and *NtHY5^CR^* seedlings through qRT-PCR. G, Quantification of nicotine content in 10-day-dark grown seedlings and seedlings transferred light for 3h in NtHY5OX and *NtHY5^CR^* seedlings through HPLC. Roots and hypocotyls were used for all the expression analysis and nicotine quantification, respectively. Tubulin was used as the endogenous control to normalize the relative expression levels. The statistical analysis was performed using two-tailed Student’s t-tests. The data are plotted as means ± SD (n=3). The error bars represent standard deviations. The asterisks indicate significant difference, *P< 0.1; **P< 0.01; ***P< 0.001.

Phytochemical analysis suggested up to 50% reduction and 75% increase in nicotine level in 20 day-old seedlings in *NtHY5^CR^* and NtHY5OX lines, respectively, compared to WT (**Figure 4D**). The levels of flavonols (kaempferol, quercetin, rutin and CGA) and anthocyanin in mutant and overexpression seedlings also got modulated (**Supplemental Figure S10A-D**; **Supplemental Figure S11A and B**) in *NtHY5^CR^* and NtHY5OX seedlings compared to WT. Thus, the decline in nicotine and flavonoid content in mutant lines and the increase in overexpression lines suggest an involvement of the NtHY5 in the biosynthesis of these molecules in tobacco seedlings.

### NtHY5 regulates light-dependent expression of genes involved in nicotine biosynthesis

In order to determine the involvement of light on the expression of genes involved in nicotine and flavonoid biosynthesis, *NtHY5^CR^* and NtHY5OX seedlings were grown in the continuous dark for 10 days, followed by exposure to light for 3h. The expression analysis suggested a significantly increased gene expression after 3h exposure to light to WT and NtHY5OX seedlings (**Figure 4E and F**). However, such an increase in the expression was not observed in *NtHY5^CR^* seedlings. We also analysed the expression of NtHY5 and flavonoid pathway genes and results suggested that exposure to 3h of light can induce the expression significantly in NtHY5OX but not in *NtHY5^CR^*plants (**Supplemental Figure S12A and B**). This induction led to enhanced accumulation of nicotine in the plants **(Figure 4G)**. These observations suggested NtHY5-dependent light responsiveness of genes involved in nicotine and flavonoid biosynthesis in tobacco.

### NtHY5 of shoot regulates nicotine biosynthesis in root

To determine light- and NtHY5-dependent regulation of nicotine biosynthesis in roots (which are not usually exposed to light), grafting experiment using shoot (scion) and root (rootstocks) of 7 day-old WT, NtHY5OX and *NtHY5^CR^* seedlings in different combinations were carried out. The grafting experiments included NtHY5OX/*NTHY5^CR^* (NtHY5OX shoot grafted over *NtHY5^CR^* root), *NtHY5^CR^*/HY5OX (*NtHY5^CR^* shoot grafted over NtHY5OX root), NtHY5OX/NtHY5OX (NtHY5OX shoot grafted over NtHY5OX root), *NtHY5^CR^*/*NtHY5^CR^*(*NtHY5^CR^* shoot grafted over *NtHY5^CR^* root), NtHY5OX/WT (NtHY5OX shoot grafted over WT root), *NtHY5^CR^*/WT (*NtHY5^CR^* shoot grafted over WT root) and WT/WT (WT shoot grafted over WT root). After three days of grafting, the roots of seedlings were covered to grow them in the dark, followed by sampling of roots after two weeks (**Supplemental Figure S13A**). The expression analysis of genes and nicotine content were measured in two ways; one in which HY5OX/HY5OX were taken as control and another with WT/WT as control.

Expression analysis of NtHY5 (**Supplemental Figure S13B**) and nicotine pathway genes for different grafting combinations suggested a significantly higher expression when NtHY5OX was used as a scion (NtHY5OX/*NtHY5^CR^*). The highest expression was observed when NtHY5OX was used for both scion and rootstock (NtHY5OX/NtHY5OX) and the lowest when *NtHY5^CR^* were used as both scion and rootstock (*NtHY5^CR^/NtHY5^CR^*) (**Figure 5A**). Similar to the expression pattern, the accumulation of nicotine followed the pattern with the highest accumulation for NtHY5OX/NtHY5OX grafted union followed by HY5OX/*HY5^CR^,* whereas the lowest for *NtHY5^CR^*/*NtHY5^CR^* grafted union followed by *NtHY5^CR^*/NtHY5OX (**Figure 5B**). Additionally, when WT/WT was taken as control, *NtHY5^CR^*/WT showed the lesser expression of NtHY5 (**Figure 5C**) and nicotine pathway genes (**Figure 5D**) compared to NtHY5OX/WT. Similar to the expression pattern, nicotine accumulation was also less in *NtHY5^CR^*/WT compared to NtHY5OX/WT (**Figure 5E**). These observations suggest that shoot NtHY5 may activate the root local HY5 which in turn binds to promoter of nicotine pathway genes. Alternatively, shoot HY5 may directly regulate the expression of nicotine pathway genes and nicotine biosynthesis. Similar to our study, previous studies have also demonstrated the movement of AtHY5 from shoot-to-root to regulate itself and other targets (Chen et al., 2016; Sharma et al., 2022).

**Figure 5.**
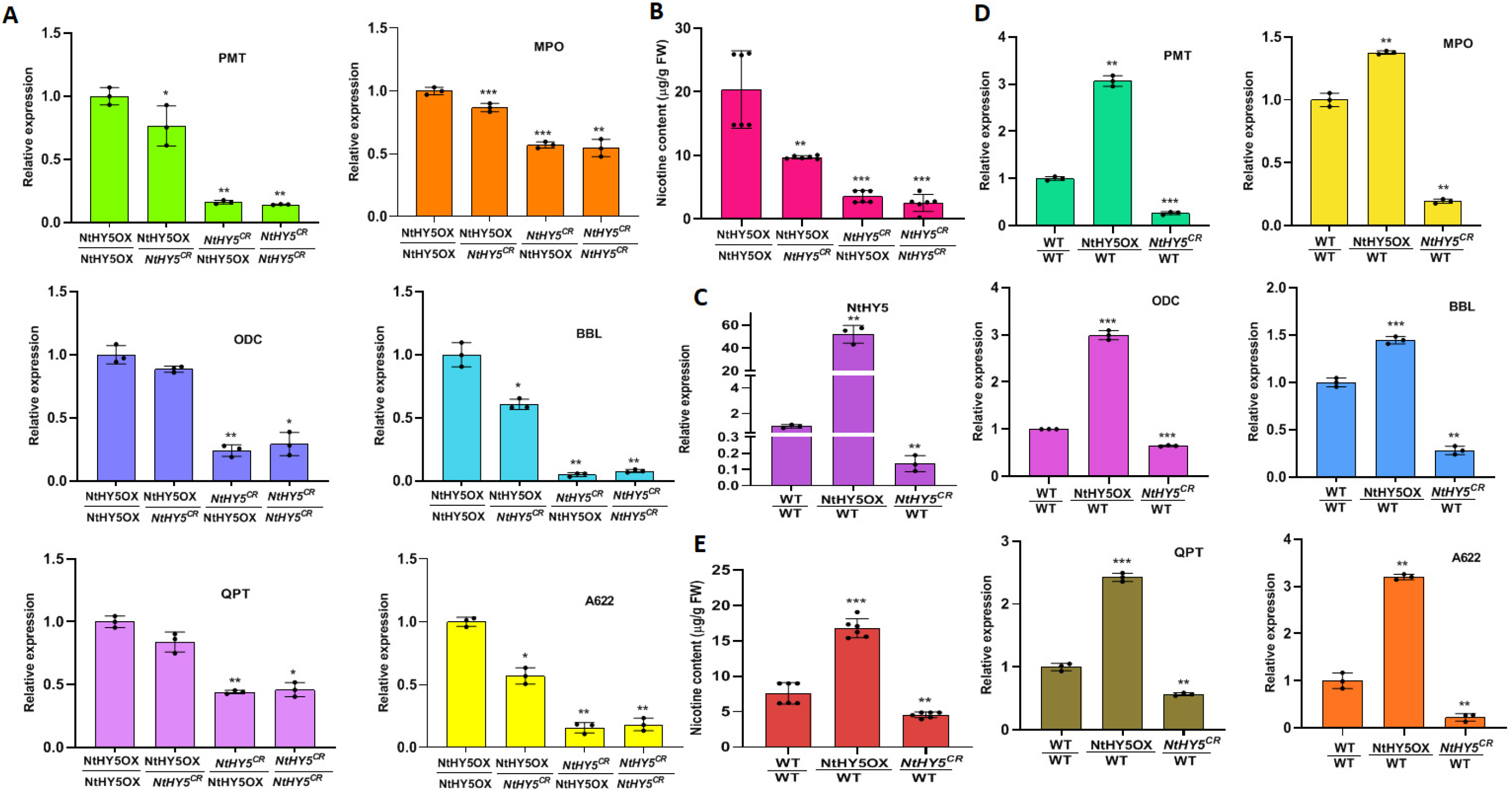
NtHY5 moves from shoot to root to modulate nicotine levels in tobacco. A, Relative expression of nicotine pathway genes (*NtODC, NtPMT, NtQPT, NtMPO, NtBBL and NtA622*) in roots of grafted NtHY5OX and *NtHY5^CR^* seedlings in several combinations keeping *HY5OX/HY5OX* as control. B, Quantitative estimation of nicotine content in grafted seedlings of WT, NtHY5OX and *NtHY5^CR^* lines. Epicotyl tissue (300 mg fresh weight) was used for the analysis. C, Relative expression of NtHY5 in roots of grafted WT, NtHY5OX and *NtHY5^CR^* seedlings in several combinations keeping *WT/WT* as control. D, Relative expression of nicotine pathway genes (*NtODC, NtPMT, NtQPT, NtMPO, NtBBL and NtA622*) in roots of grafted WT, NtHY5OX and *NtHY5^CR^* seedlings in several combinations keeping *WT/WT* as control. E, Quantitative estimation nicotine content in grafted seedlings of WT, NtHY5OX and *NtHY5^CR^* lines keeping *WT/WT* as control using HPLC. Roots and hypocotyls were used for all the expression analysis and nicotine quantification, respectively. TUBULIN was used as the endogenous control to normalize the relative expression levels. The statistical analysis was performed using two-tailed Student’s t-tests. The data are plotted as means ± SD (n= 3). The error bars represent standard deviations. The asterisks indicate significant difference, *P< 0.1; **P< 0.01; ***P< 0.001.

### NtHY5 regulates nicotine biosynthesis in mature tobacco plants

Our abovementioned results using NtHY5OX and *NtHY5^CR^*seedlings suggested that NtHY5 regulates nicotine biosynthesis in tobacco roots. We explored the impact of NtHY5 on nicotine synthesis in mature plants. The expression analysis suggested increased expression of genes involved in nicotine biosynthesis in NtHY5OX compared to *NtHY5^CR^* and WT roots (**Figure 6A and B**). At the metabolite level, leaves of NtHY5OX accumulated significantly higher nicotine levels compared to WT and *NtHY5^CR^* leaves. There was 92% and 70% reduction in nicotine content in *NtHY5^CR^* leaves compared to NtHY5OX and WT leaves, respectively (**Figure 6C**).

**Figure 6.**
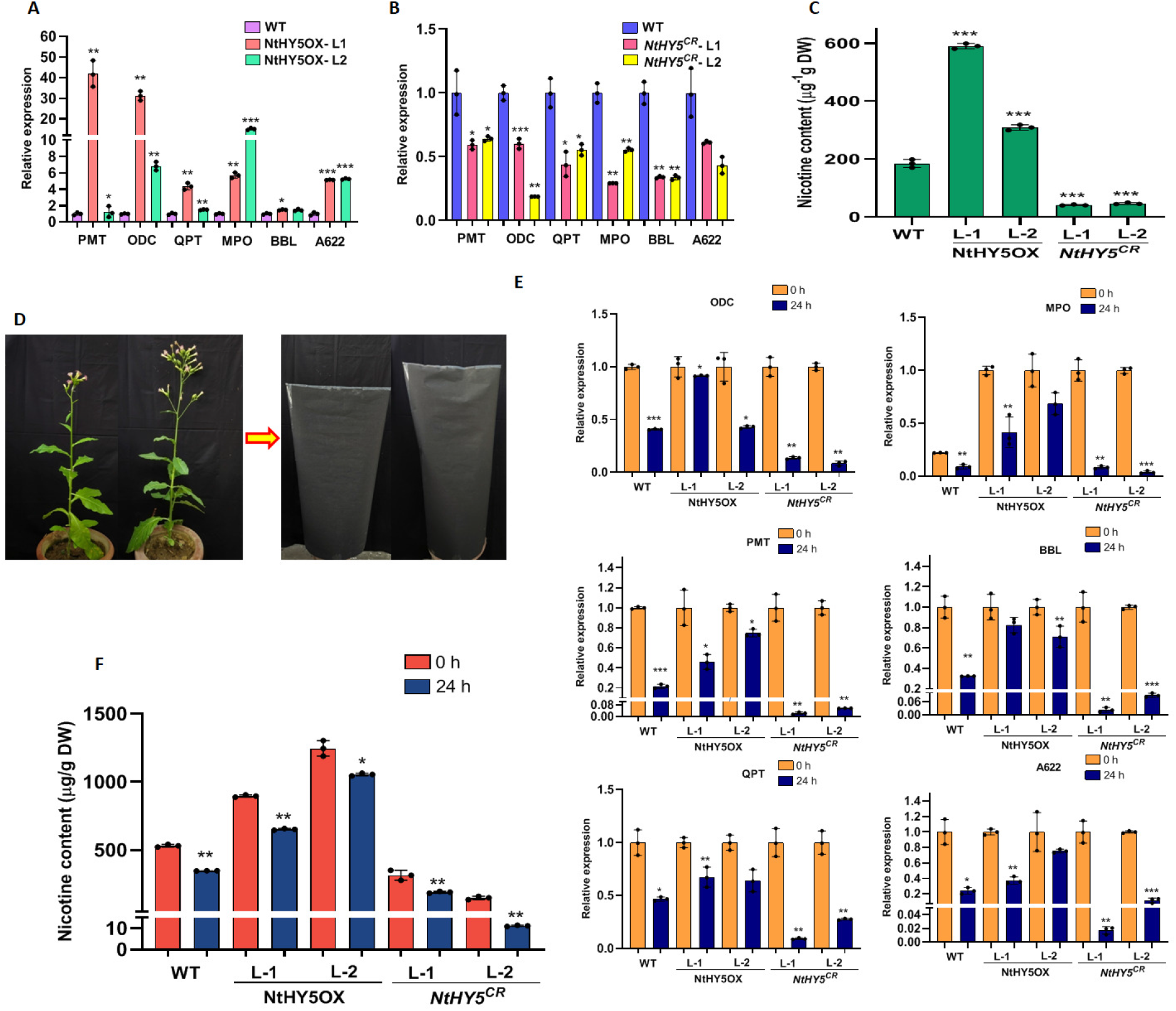
NtHY5 modulates nicotine levels in mature tobacco transgenic lines. A and B Relative transcript abundance of nicotine pathway genes (*NtODC, NtPMT, NtMPO, NtQPT and NtA622*) in roots of NtHY5OX and *NtHY5^CR^* 3-month-old tobacco plants. C, Quantitative estimation nicotine content in 3-month-old leaves of plant using HPLC. D, Representative photograph of 5-months-old mature tobacco plant being kept in dark for 24hours. E, Relative expression analysis of nicotine pathway genes (*NtODC, NtMPO, NtPMT, NtA622, NtQPT and NtBBL*) in roots of 5-months-old mature NtHY5OX and *NtHY5^CR^* plants grown in light and transferred to dark for 24h through qRT-PCR. F, Quantification of nicotine content in in 5-months-old mature tobacco leaves transferred to dark for 24h in NtHY5OX and *NtHY5^CR^* lines through HPLC. Leaf tissue (300 mg dry weight) was used for the analysis. Roots and hypocotyls were used for all the expression analysis and nicotine quantification, respectively. Tubulin was used as the endogenous control to normalize the relative expression levels. The statistical analysis was performed using two tailed Student’s t-tests. The data are plotted as means ± SD (n= 3). For measurement of root bending angle, root length, hypocotyls length and seed size (n=15). The error bar represents standard deviations. The asterisks indicate significant difference, *P< 0.1; **P< 0.01; ***P< 0.001.

We also analyzed the effect of light-dark transition on nicotine biosynthesis using NtHY5OX and *NtHY5^CR^* plants. Light-grown mature tobacco plants were completely covered with black paper for 24 h before sampling for the analysis **(Figure 6D)**. Roots of both NtHY5OX and *NtHY5^CR^* plants showed reduced expression of nicotine pathway genes during the light-to-dark transition. However, the reduction was substantially higher in *NtHY5^CR^* plants than in NtHY5OX when compared to WT (**Figure 6E**). The modulation in the expression of genes led us to determine nicotine concentration in mature leaves. The analysis suggested a higher accumulation of nicotine in NtHY5OX leaves than in *NtHY5^CR^* compared to WT after the light-to-dark transition **(Figure 6F)**. This suggested that dark negatively regulated nicotine synthesis in mature tobacco plants.

### NtHY5 regulates growth and development of tobacco

Previous study suggested that flavonols can also function as an endogenous polar auxin transport inhibitor to regulate shoot and root growth under various conditions (**Peer et al., 2004, Santelia et al., 2008**) although the relation between hormone signaling and NtHY5 could be the subject of further study and reduction in nicotine content leads to positive regulation of root growth (**Hildreth et al., 2011**). We subsequently studied the effect of modulated nicotine and flavonoids on the growth and development of tobacco. A comparative analysis between NtHY5OX and *NtHY5^CR^*transgenic lines suggested differences in the growth of NtHY5OX and *NtHY5^CR^*light-grown seedlings (**Supplemental Figure S14A**). Several previous studies have demonstrated that AtHY5 regulates the hypocotyl length by repressing the expression of hypocotyl cell elongation-related genes (Liu et al., 2013). To validate the role of NtHY5 in hypocotyl length elongation in tobacco, we studied the difference in hypocotyl length in NtHY5OX and *NtHY5^CR^*plants. The analysis suggested that the hypocotyl length of *NtHY5^CR^*mutant seedlings was significantly higher compared to overexpression lines (**Supplemental Figure S14B**). At the same time, the hypocotyl length of the dark-grown overexpression lines was significantly higher compared to *NtHY5^CR^* mutant plants (**Supplemental Figure S14C and D**). We also measured the root length and bending angle of 10-d-old light-grown seedlings. Analysis suggested the higher root length of *NtHY5^CR^* seedlings compared to WT and NtHY5OX, whereas the root bending angle of NtHY5OX seedlings was higher compared to WT and *NtHY5^CR^* plants (**Supplemental Figure S14E** and **F**).

We also observed the modulation in seed size; *NtHY5^CR^*plants produced smaller seeds (**Supplemental Figure S15 A and B**) compared to NtHY5OX and WT (**Supplemental Figure S15C**). In addition to these factors, we observed the difference in height and chlorophyll content of the young plants. The plant height of 40-days-old *NtHY5^CR^* plants was significantly higher than *NtHY5* overexpression and WT plants (**Supplemental Figure S15D and E**), whereas chlorophyll was found to be lesser in *NtHY5^CR^* plants compared to NtHY5OX and WT (**Supplemental Figure S15F**).

### NtHY5 enhances the salt stress tolerance in tobacco

As HY5 is known to play a role in abiotic stress tolerance, the response of NtHY5 under salt stress was analyzed through leaf disc assay using NaCl (200 mM), and NtHY5OX, *NtHY5^CR^* and WT plants. After 3 days of salt stress, higher chlorosis was observed in *NtHY5^CR^* leaves compared to leaves of NtHY5OX and WT plants (**Supplemental Figure S16A and B**). Chlorosis affects the survival of leaves by decreasing the chlorophyll content (Chen et al., 2019). Thus, the survival of leaves was measured by analyzing the total chlorophyll content as well as chlorophyll a and b. A higher chlorophyll content was observed in NtHY5OX leaves compared with *NtHY5^CR^* and WT leaves after 3 days of salt stress (**Supplemental Figure S16C**). It is well known that salt stress leads to excessive accumulation of ROS, leading to cell death (Chen et al., 2019). Higher NBT and DAB staining were observed in *NtHY5^CR^* compared with NtHY5OX and WT (**Supplemental Figure S16D and E**), respectively, suggesting the higher ROS formation in *NtHY5^CR^* leaves, thus above study concluded that NtHY5 might play a crucial role in salt stress tolerance in tobacco by inhibiting ROS accumulation. A higher ROS accumulation results in lower antioxidant activity in *NtHY5^CR^*lines, whereas lower ROS accumulation leads to the higher antioxidant activity in NtHY5OX lines compared to WT (**Supplemental Figure S17**).

## Discussion

The nicotine biosynthesis pathway in terms of intermediate steps and enzymes is well explored (Dewey and Xie, 2013). Additionally, several previous studies suggested the role of various transcription factors belonging to different families, for example, helix–loop–helix (bHLH) transcription factors (e.g. MYC2a/b), MYC2-type bHLH TFs NbbHLH1 and NbbHLH2, AP2/ERF genes, NtORC1/ERF221 and NtJAP1/ERF10, and MYB (NtMYB305a), in regulating nicotine biosynthesis (Song et al., 2011; Shoji et al., 2011; Shear et al., 2014; Liu et al., 2019; Bian et al., 2021). In the past, other factors, including insect attack, wounding, and jasmonate (JA) have been shown to be involved in the biosynthesis of nicotine (Imanishi et al., 1998; Shi et al., 2006; Bozorov et al., 2017; Li et al., 2018; Liu et al., 2020). Previous study suggested the presence of several G and GCC boxes in the promoter regions of nicotine biosynthesis genes (Zhang et al., 2011). It has also been established that HY5 binds to G-box (CACGTG) in the promoters of light responsive genes. Therefore, we hypothesized that NtHY5 could regulate the nicotine production in tobacco.

Previous studies in *Arabidopsis* and other plant species suggested HY5 as a master regulator of photomorphogenesis and a key signaling component of light. It promotes photomorphogenesis under a broad spectrum of wavelengths (Ulm et al., 2004; Lee et al., 2007; Nawkar et al., 2017). However, the functional characterization of tobacco HY5 is not done yet. In this study, we have identified the NtHY5 and demonstrated that it regulates nicotine biosynthesis in tobacco, extending our understanding of NtHY5-mediated regulation of secondary plant metabolism. The present work shows the involvement of light signaling component in nicotine and flavonoid regulation of tobacco as accumulation of nicotine as well as flavonols also found to be higher in light-grown seedlings compared to dark-grown seedlings (**Figure 1D and Supplemental Figure S1B**). WT dark-grown seedlings showed the induction in the expression of nicotine pathway genes on exposure to light and continuously increased with increase in time interval till saturation (**Figure 1E**). Thus, above results led us to hypothesize the involvement of NtHY5 as one of the potential factors responsible for light-mediated nicotine and flavonoid regulation in tobacco. A higher expression of identified NtHY5 in light-grown seedlings as well as our further analysis in which we have exposed 10-d-old dark-grown seedlings to light of different time interval suggested the significant induction in expression of nicotine pathway genes (**Figure 1E and F**) as well as indicated, NtHY5 as light-responsive key regulatory factor in nicotine and flavonoid biosynthesis (**Supplemental Figure S1**) in tobacco.

Prior studies have established that HY5 acts as a transcription factor that predominantly binds to the LREs like *ACGT*-containing *cis*-element (e.g., *G-box* and *T/G-box*) and controls the expression of numerous target genes in response to light signals (Song et al., 2008). To investigate the binding of NtHY5 to the promoter of nicotine pathway genes, *in silico* analysis was carried out and the result revealed the presence of cis-acting LREs upstream of the transcription start site on promoters of PMT, QPT, ODC and NtMYB12. Similar to our result, there are previous studies, suggesting the presence of GCC and G-box elements in the promotor regions of nicotine pathway genes (Todd et al., 2010; Shoji and Hashimoto, 2011; Zhang et al., 2012). Our EMSA analysis validated NtHY5 intraction with the promoters of genes involved in nicotine pathway and flavonoid regulatory transcription factor, suggesting the involvement of NtHY5 in nicotine and flavonoid biosynthesis (**Figure 2 and Supplemental Figure S6**).

Previously, several HY5 orthologs from different plant species have been identified and characterized (Nishimura et al., 2002; Burman et al., 2018). OsZIP48 is reported as a functional ortholog of AtHY5 in rice as it was able to rescue the phenotype of *Arabidopsis hy5* mutant plants after complementing with OsZIP48. Similarly, overexpression of STF1, the HY5 ortholog in soybean, can fully complement the *hy5* phenotype (Song et al., 2008). As NtHY5 was identified for the first time in the current study thus complementation study has been done to identify whether NtHY5 can fully complement the *Arabidopsis hy5* or not? The current study demonstrated that NtHY5 partially complemented *Arabidopsis hy5* mutant for phenotypes such as hypocotyl length, root length, seed size and rosette diameter. This study also indicated the fully complementation at the level of expression of structural and regulatory genes of flavonoid biosynthesis and accumulation of metabolites, such as flavonoid, anthocyanin, lignin and chlorophyll in *NtHY5; hy5* compared to *hy5* **(Figure 3 and Supplemental Figure S7)**. These findings suggest that NtHY5 had a conserved function across different species, and it regulates flavonoid biosynthesis in complemented lines of *Arabidopsis*.

To decipher the molecular mechanism through which light regulate nicotine and flavonoid biosynthesis in tobacco, CRISPR/Cas9-mediated knockout and overexpression tobacco plants for NtHY5 were developed. As per previous studies, nicotine synthesis takes place in the root, which is subsequently transported to the leaf via the xylem (Hayashi et al., 2020). Thus, in the current study, analysis of nicotine content and pathway gene expression was carried out at seedling (roots were kept in the dark) as well as mature plant stage in mutant and overexpressed plants of NtHY5 compared to WT. The expression analysis of seedlings as well as roots of mature plants indicated a higher expression of nicotine pathway genes in seedlings and roots of NtHY5OX mature plants, whereas expression was lower in seedlings and roots of *NtHY5^CR^*plants in comparison to WT (**Figure 4C, 6A and 6B**). Quantitative estimation of nicotine content in seedlings suggested up to 50% reduction and four-fold increase in nicotine level in *NtHY5^CR^* and NtHY5OX lines, respectively. At the mature plant stage, leaves of NtHY5OX accumulated 92% higher nicotine levels compared to WT and *NtHY5^CR^* leaves showed 70% reduction in nicotine content in comparison to WT. Thus, current study reveals that NtHY5 regulates nicotine synthesis not only at seedling but also at the mature plant stage. The CRISPR/cas9 mutated NtHY5 could not bind to promoters of nicotine pathway genes, resulting into a lower accumulation of nicotine. The current study also indicated that NtHY5 is induced by light supported by dark-to-light induction analysis. In OE lines, NtHY5 regulates its own expression in light, while in dark it undergoes degradation and is unable to regulate its own expression. This leads to a lower expression in the dark compared to seedlings exposed to 3h light after 10-day growth in dark **(Supplemental Figure S12 A)**. Less nicotine-containing edited tobacco plants will be beneficial for mankind by helping adults to switch instead of continuing smoking cigarettes because nicotine is part of what makes cigarettes addictive.

As tobacco roots are known to synthesize nicotine (Hayashi et al., 2020) and are typically in the dark, we studied how shoot NtHY5 regulates nicotine production in roots. We carried out grafting experiments in which several hypocotyl graft chimeras with roots covered in darkness were examined. We observed that a NtHY5OX scion facilitated an enhanced nicotine accumulation (as opposed to a *NtHY5^CR^* scion) (**Figure 5, Supplemental Figure S13**), indicating that a shoot derived and mobile NtHY5 signal controls nicotine biosynthesis in roots, controlling both by directly modulating nicotine pathway genes and automatically activating local HY5 in the roots. Similar to our findings, a previous study suggested that *Arabidopsis* HY5 regulates growth in response to light, is a shoot-to-root mobile signal that mediates light promotion of root growth and nitrate uptake (Chen et al., 2016).

Our tissue-specific expression analysis also suggested the lower expression of nicotine pathway genes in leaves compared to roots. In the past, several studies have also suggested the biosynthesis of nicotine in the cortical cells of roots through various enzymatic steps, which is further transported to leaves vacuoles via xylem (Shoji et al., 2009; Dewey et al., <u>2013</u>; Hayashi et al., 2020; Shoji et al., 2021). This was the reason for using roots for all the expression analysis whereas young and mature leaves were used for the nicotine quantitative analysis. However, it is possible that NtHY5 might regulates nicotine biosynthesis in the leaves but to a lesser extent.

In case of *NtHY5^CR^*/NtHY5OX grafted seedling, no functional HY5 is present in shoot which can move to root to activate root local HY5; thus, expression of nicotine pathway genes is lower. In addition, *NtHY5^CR^*/NtHY5OX grafted seedling, though HY5 in root is expressing under control of CaMV35S promoter, the possibility of degradation of this by root-specific COP1 cannot be ruled out. This might be one of the reasons for the low nicotine accumulation in *NtHY5^CR^*/NtHY5OX grafted seedling. NtHY5 transcript levels are very high in roots of NtHY5OX/*NtHY5^CR^*grafted seedlings. There could be a possibility that transcript levels observed are from the mutated gene or shoot NtHY5 transcripts which move to the roots. There are a few previous studies that suggest that transcripts could also move between scion and rootstock. For example, CmGAI transcripts in tomato, BEL5 transcripts in tomato and transcripts in grapes have been demonstrated to be present from scion to rootstock (Yin et al., 2022; Haywood et al., 2005; Banerjee et al., 2006; Yang et al., 2015).

We also demonstrated the regulation of flavonols and anthocyanin biosynthesis by NtHY5 as they are found to be reduced in NtHY5 knockout seedlings in comparison to WT (**Supplemental Figure S10 and S11**). We found a significant increase in flavonol content, but interestingly decrease in anthocyanin content in NtHY5OX lines, suggesting the diversion of flux towards flavonol biosynthesis. Thus, these results provided a deeper insight of light-mediated regulatory mechanism involved in nicotine and flavonoid biosynthesis in tobacco.

HY5 and specific MYBs are known to be involved in plant growth and development (Zhong et al., 2008; McCarthy et al., 2009; Gangappa et al., 2016; Misra et al., 2010). OsbZIP48 ortholog is reported to cause a reduction in cell size, thus reducing the plant height of OsbZIP48 overexpression transgenic plants (Burman et al., 2018). HY5 negatively regulates hypocotyl elongation by suppressing auxin signaling (Cluis et al., 2004; Sibout et al., 2006). Our results suggested that NtHY5 regulates flavonol content in tobacco. Similarily, previous study suggested that flavonols can function as an endogenous polar auxin transport inhibitor to regulate shoot and root growth under various conditions (Peer et al., 2004; Santelia et al., 2008). Thus, we next determined the impact of NtHY5 on plant growth parameters since, no data is available on the involvement of NtHY5 in the growth and development of tobacco via regulating secondary metabolites. We conducted a comparison study between the transgenic NtHY5OX and *NtHY5^CR^* lines. The results suggested that modulation in the NtHY5 expression causes changes in nicotine and flavonoid concentration leading to variation in various morphological characteristics, with contrasting phenotypes between NtHY5OX and *NtHY5^CR^* plants. Hypocotyls length of mutants was significantly longer than overexpressed and WT seedlings grown in light for 10 days. Interestingly, shorter hypocotyl length was observed in mutants in comparison to overexpress and WT in dark-grown seedlings (**Supplemental Figure S14 A-D**). This suggested that mutation in NtHY5 might have eradicated the light-dependent repression of cell elongation and does not promote cell elongation independent of the light stimulus.

In tobacco, one previous study suggested that RNAi silencing of NUP1, a plasma membrane localized nicotine transporter, causes a reduction in nicotine content which in turn regulates the root growth of seedlings positively (Hildreth et al., 2011). In our study, we demonstrated that low nicotine content, accompanied by low flavonols, causes an increase in root length in *NtHY5^CR^*seedlings compared to overexpression lines and WT. It is earlier reported that the asymmetric accumulation of flavonoids in the root tips of *Arabidopsis* affects the direction of root growth (Wan et al., 2018). In agreement, root bending angle of 10-day-old light-grown NtHY5 overexpression and *NtHY5^CR^* seedlings of the tobacco was measured. The result suggested the higher root bending angle in NtHY5OX lines in comparison to *NtHY5^CR^* and WT (**Supplemental Figure S14F**). Interestingly, analysis also showed the modulation in tobacco seed size and height of the 40-d-old plants (**Supplemental Figure S15B and FF**). Thus, results indicated that NtHY5 plays a pivotal role in tobacco growth and development through auxin signaling, although the relation between hormone signaling and NtHY5 could be the subject of further study.

Our study suggested the regulation of NtMYB12 by the NtHY5 through interaction with its promoter. One recent study demonstrated that overexpression of NtMYB12 causes more flavonol accumulation and a higher ROS scavenging property, enhancing tobacco tolerance to low Pi stress (Song et al., 2020). There are some studies which suggest that tobacco responds to salt stress by increased activity of antioxidant enzymes (Sun et al., 2020. In our study, antioxidant activity was measured, and the analysis suggested higher activity in overexpression lines compared to WT and *NtHY5^CR^* plants. The high antioxidant activity led to lesser chlorosis in salt-treated young leaves of NtHY5OX lines compared with *NtHY5^CR^* **(Supplemental Figure S16A-C**), suggesting that higher NtHY5 expression leads to higher ROS scavenging ability in overexpression lines. As tobacco *NtHY5^CR^* mutants were salt-sensitive, thus more ROS accumulated in salt-treated mutant leaves **(Supplemental Figure S16D and E**). Thus, our study suggests that NtHY5 promotes salt stress tolerance at least partially through the antioxidant activity of flavonoids **(Supplemental Figure S17)**. To understand whether nicotine also has the antioxidant activity, a detailed regulatory mechanism needs to be studied.

We propose the model (**Figure 7**), which depicts that HY5 is a shoot-root mobile signal that mediates the light-regulated synthesis of nicotine and flavonoids. This coupling is achieved via the regulation of nicotine and flavonoid pathway genes through interaction of the NtHY5 with the G Boxes present within the promoters of different genes. NtHY5 is also found to modulate growth and development, as evident through modulated seed size, plant height and root length. Additionally, NtHY5 provides salt tolerance via diminishing ROS accumulation evident through DAB and NBT staining. Thus, this study provides an essential insight into a molecular strategy to develop tobacco having significantly low nicotine through manipulating the NtHY5 gene of tobacco.

**Figure 7.**
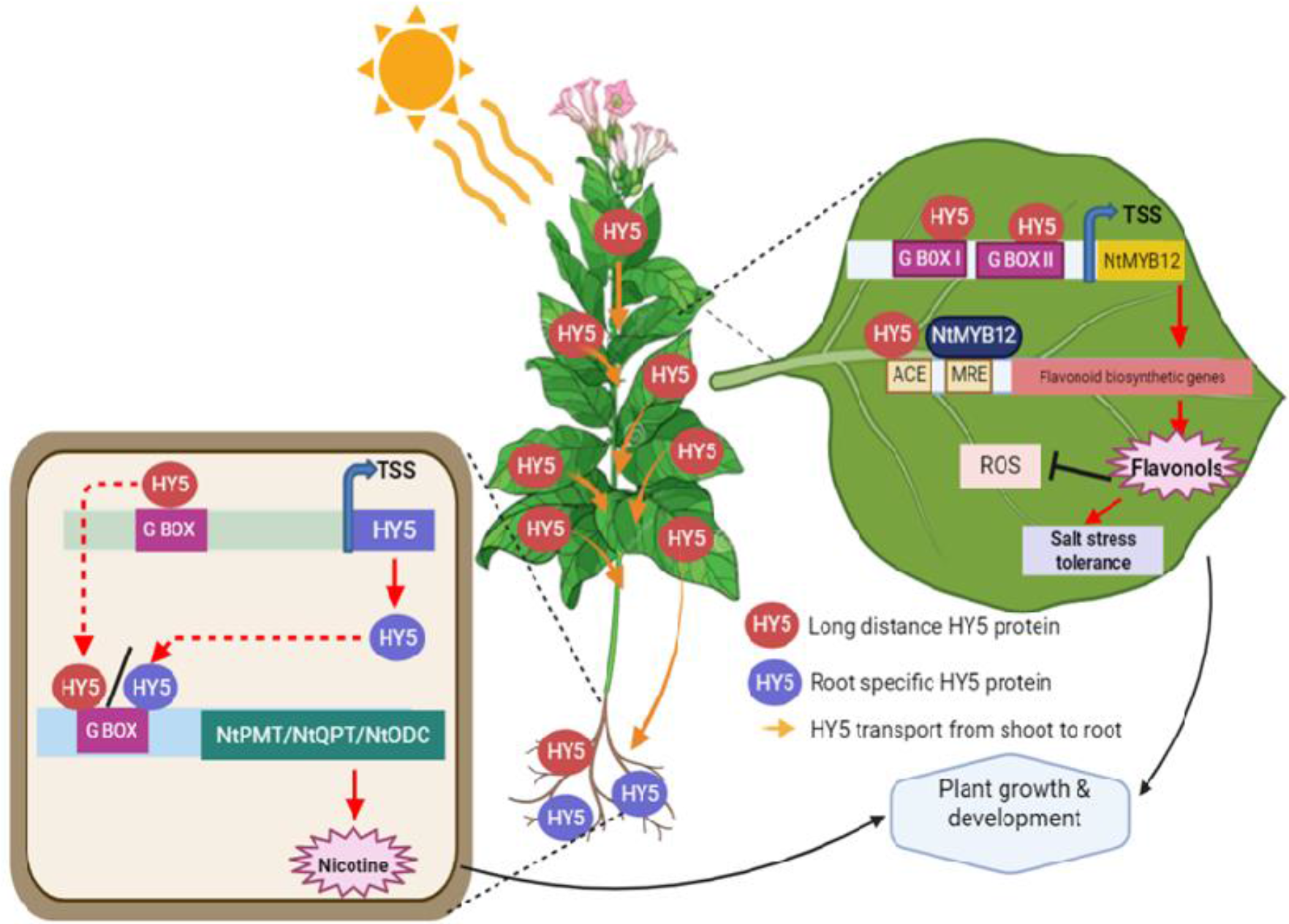
Proposed working model showing the role of NtHY5 in nicotine and flavonol accumulation in tobacco. We propose that NtHY5 moves from shoot to root to upregulate the expression of nicotine pathway genes in light which results into increased accumulation of nicotine as well as flavonoid biosynthesis. NtHY5 regulates nicotine and flavonol biosynthesis via binding to the G-boxes in the promoters of nicotine and flavonol regulatory genes. Higher accumulation of secondary metabolites leads to less ROS accumulation thus increased salt stress tolerance ability. Higher nicotine and flavonoids also regulates growth and development of tobacco. Arrowheads and tees indicate positive and negative regulation, respectively.

## Materials and Methods

### Isolation and sequence analysis of NtHY5

To identify the ortholog of AtHY5 in tobacco, BLAST searches were performed on National Center for Biotechnology Information (http://www.ncbi.nlm.nih.gov/BLAST). The sequence was retrieved and amino acid sequence was analyzed for the leucine rich bZIP Zinc finger domain and VPE domain using multallignment tool. Conserved domain database (CCD) was employed to ensure the bZIP family of the identified NtHY5 protein. Promoter sequences (∼1.5 kb upstream of translation start site) of NtMYB12 gene was retrieved from the respective databases AGRIS (Palaniswamy et al., 2006) and PlantCare (Lescot et al., 2002), which was utilized for the scanning of light-responsive cis-elements (LREs). Homologs of AtHY5 that belongs to different families were identified by using the National Center for Biotechnology Information BLASTn tool. The genes retrieved were checked for the presence of the requisite domains using InterProScan (Quevillon et al., 2005) (http://www.ebi.ac.uk/Tools/pfa/iprscan/). A phylogenetic tree was constructed using the Neighbor Joining (NJ) algorithm in MEGA6.0 software (Kumar et al., 2008).

### Plant material, growth conditions and treatments

Leaves of *Nicotiana tabacum* cv. *Petit Havana* (NtPH) have been used for raising the transgenic plants. Tobacco plants were grown in glass house at 25°C±2°C and 16 h/8 h light-dark photoperiods for harvesting of tissues belonging to different developmental stage. Seeds were surface sterilized using 70% EtOH for 1 min and then dipped in 50% bleach for 10 min, washed repeatedly with autoclaved Milli-Q water and grown with roots covered in dark for 20 days. For dark treatment, seeds were surface sterilized and grown for 10 days in absence of light, 24°C temperature, and 60% relative humidity on one-half-strength Murashige and Skoog (MS) medium (Sigma) plates after stratification for 2 to 3 d at 4°C in the dark and subsequently transferred to culture room with 16 h/8 h light-dark photoperiods and 150– 180 mm m^-2^s ^-1^ light intensity. For light-to-dark treatment glass house grown 5-month old tobacco plants were completely covered with black paper for 24h before sampling. In mature plants, three biological replicates of each line were grown in glasshouse till maturity. Plants were then uprooted and the roots were cut from the root-shoot junction and sampled for further analysis. In seedlings, seeds were surface sterilized and grown with roots covered in dark for 20 days following which the roots were cut from root-shoot junction in dark room and sampled for further analysis.

*Arabidopsis* (Col-0) was used as the wild-type plant in complementation study. Seeds were surface sterilized and after stratification transferred to a growth chamber under controlled conditions of 16-h-light/8-h-dark photoperiod cycle, 22°C temperature, 150 to 180 µmol m^-2^s^-1^ light intensity, and 60% relative humidity for 10 days unless mentioned otherwise.

### Plasmid construction and generation of *Arabidopsis* and tobacco transgenic and mutant plants

To overexpress NtHY5, the NtHY5 was amplified from the single-stranded complementary DNA (cDNA) library prepared from 10-day-old tobacco seedlings through PCR using specific primers. The full-length open reading frame of the NtHY5 cDNA, under the control of cauliflower mosaic virus 35S promoter in the binary vector pBI121 (Clontech, USA) was transferred into *Agrobacterium tumefaciens* strain GV3101 and used to transform *Arabidopsis* by floral dip method and tobacco plants by leaf disk method (Horsch et al., 1985). Several transgenic tobacco lines constitutively expressing NtHY5 cDNA were selected on the basis of RT-PCR. Seeds were harvested, sterilized and plated on solid half strength MS medium supplemented with 100 mg/l kanamycin. Antibiotic resistant plants were shifted to the glass house and grown until maturity.

For generation of mutant plants of NtHY5, CRISPR/Cas9 technology was used. A 20-bp gRNA was selected from the identified coding region. NtHY5 gRNAs was cloned into the binary vector pHSE401 using the BsaI restriction site (Xing et al., 2014). This vector contains Cas9 endonuclease-encoding gene under dual CaMV35S promoter as well as genes encoding neomycin phosphotransferase and hygromycin phosphotransferase as selection markers. All the constructs were sequenced from both the orientations using plasmid-specific forward and vector reverse primers after that transferred to GV3101 and then transformed to tobacco.

### Gene expression analysis

Total RNA was isolated using a Spectrum Plant Total RNA kit (Sigma Aldrich) following the manufacturer’s instruction. For qRT-PCR analysis, 2 µg of DNA free RNA was reverse transcribed using a RevertAid H minus first-strand cDNA synthesis Kit (Fermentas) according to the manufacturer’s instructions. qRT-PCR was performed using Fast Syber Green mix (Applied Biosystems) in a Fast 7500 Thermal Cycler instrument (Applied Biosystems). Expression was normalized using Tubulin (NtTUB, XM_009622503; AtTUB, NM_125664) and analyzed through the comparative CT method (Livak and Schmittgen 2001). For each experiment, three independent RNA preparations and total nine technical replicates (three for each RNA preprations) were used. The oligonucleotide primers used to study expression of different genes were designed using the Primer Express 3.0.1 tool (Applied Biosystems) and information is provided in Supplementary Table S5.

### Tobacco grafting

Grafting between wild type, NtHY5OX, *NtHY5^CR^* lines was performed according to method described by Rus et al. (2006). Seedlings grown on one-half-strength MS media for 7 days were used for grafting. Cuts between the scion and stock were made at an angle of 90° and were then placed in desired combinations. The grafted seedlings remained on the media plate for a period of 3 days to allow the formation of the graft union. Roots of the grafted seedlings were further covered in darkness for 2 more weeks prior to sampling. For further experiments, only the grafted seedlings were used. Seedlings which did not produce proper graft (scion and rootstock did not bind with each other and rootstock ultimately died) were discarded.

### Extraction and quantitative estimation of nicotine

Nicotine extraction was carried out with slight modifications (Khan et al., 2017). About 300 mg freeze-dried tissue samples for mature leaves and 300 mg fresh weight tissue for seedlings/ grafted seedlings were grinded to powder and extracted in 2.0 mL 75 % methanol. The extract was ultrasonicated for 60 min, and then centrifuged (14,000 rpm) for 10 min. The supernatant was concentrated to dry powder and then dissolved in 1mL HPLC grade methanol. Extracts were filtered through 0.2 mm filter (Millipore, USA) and subjected to HPLC analysis. Lab Solutions software (Shimadzu) was used for the quantification of nicotine through HPLC. Column used was Phenomenex Luna C18(2) column (250 mm x 4.6 mm x 5µ). Mobile phase consisted of water: methanol: 0.1 M buffer acetate (pH4.5): acetonitrile: acetic acid in the ratio 74: 3: 20: 2:1 (Solvent A) with pH adjusted to 4.2 with triethylamine. Solvent A was maintained at 100% throughout the run with a slight modification in the gradient of flow rate of 1ml/min in 0-15min, 1.0-1.5ml/min in 15-15.5min, 1.5ml/min in 15.5-24.5min, 1.5-1.0ml/min in 24.5-25min and 1ml/min in 25-30min. The samples were analysed by HPLC–PDA and chromatograms were recorded at 254nm (Pereira et al., 2001).

### Extraction and quantitative estimation of flavonols

Extraction of flavonols was carried out by grinding the plant material (300mg) into the fine powder in liquid N_2_ and then extracted with 80% methanol overnight at room temperature with brief agitation. The extract was hydrolyzed with an equal amount of 6 N HCl at 70°C for 40 min followed by addition of an equal amount of methanol to prevent the precipitation of the aglycones (Jiang et al. 2015). Extracts were filtered through 0.2 mm filter (Millipore, USA) before HPLC. For non-hydrolyzed extracts, samples were extracted as described previously (Pandey et al., 2014). The mobile phase was a gradient prepared from 0.05% (v/v) ortho-phosphoric acid in HPLC-grade water (component A) and methanol (component B). Before use, the components were filtered through 0.45-mm nylon filters and de-aerated in an ultrasonic bath. The gradient from 25 to 50% B in 0–3 min, 50 to 80% B in 3–18 min, 80 to 25% B in 25 min and 25% B in 30 min was used for conditioning of the column with a flow rate of 1 ml/min. All the samples were analysed by HPLC–PDA with a Waters 1525 Binary HPLC Pump system comprising PDA detector (Niranjan et al., 2011).

### Extraction of total anthocyanin, chlorophyll and lignin staining

For anthocyanin, 300 mg tissue were boiled for 3 min in extraction solution (propanol: HCl: H2O, 18:1:81), and incubated at room temperature in the dark for at least 2 h. Samples were centrifuged, and the absorbance of the supernatants was measured at 535 and 650 nm. Anthocyanin content was calculated as

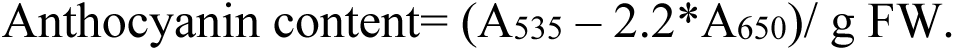

Chlorophyll content was measured by using the 300mg tissue of 10-d-old WT and transgenic seedlings. Sample was crushed in 2 ml (80 %, v/v) chilled acetone. Homogenized tissues were centrifuged at 7,800g for 10 min. Supernatant was collected and absorbance was recorded at 663, 645, 510 and 480 nm. Total chlorophyll content was calculated in mg g^−1^ fresh wt using formula (Machlachalan and Zalik, 1963).

To visualize lignified cells in stems, hand-cut sections were stained using phloroglucinol (Sigma Aldrich) for 1 min and visualized on a Leica DM2500 microscope (Sharma et al., 2020).To measure the length of vascular bundle, automated length measuring scale of Leica DM2500 was used.

### Determination of total antioxidant activity

A method with slight modification was used to determine the antioxidant activity. DPPH (1, 1-diphenyl-2-picrylhydrazyl free radical) assay was used in determining the free radical scavenging activity (Wong et al., 2006). In brief, 0.1 mM solution of DPPH was prepared in methanol. The initial absorbance of DPPH in methanol was measured at 515 nm and did not change throughout the period of assay. An aliquot of the samples (50–100 ml) was used to make up volume to 1.5 ml with DPPH solution. The sample was incubated at room temperature for 30 min. The absorbance of the samples was recorded at 515 nm using Ultraspec 3000 UV/Vis spectrophotometer (Pharmacia Biotech Ltd., Cambridge, CB4, 4FJ, UK). Trolox was used to make the calibration curve using different dilutions. The antioxidant capacity based on the DPPH free radical scavenging ability of the extract was expressed as mM Trolox equivalents per gm of plant material on fresh weight basis.

### Expression and purification of NtHY5

The complete open reading frame of NtHY5 was amplified and cloned into pET-28b(+) vector (Novagen, Germany). The construct was transformed into *E. coli* BL21 (DE3) pLysS (Invitrogen, USA) for prokaryotic expression, induced by IPTG. Recombinant 6X-Histagged HY5 was purified using a Ni-NTA column (Nucleopore, India) and the eluted and was quantified through Bradford assay.

### Electrophoretic mobility shift assay.

The 2^nd^ generation DIG Gel Shift EMSA kit (Roche, USA) was used to label the probes with digoxigenin as per manufacturer’s instructions. Labelled probes were incubated at 21°C for 30 min in binding buffer [100 mM HEPES (pH 7.6), 5 mM EDTA, 50 mM (NH4)_2_SO_4_, 5 mM DTT, Tween 20, 1% (w/v), 150 mM KCl] with or without recombinant protein (500 ng). Unlabeled probes were added to the reaction solution in increasing concentrations to test specific binding. The binding reaction was resolved on a 6% polyacrylamide gel in 0.5X TBE (pH 8.0) buffer and was semi-dry blotted (Transblot, BIO-RAD, USA) onto a positively charged nylon membrane (BrightStar, Invitrogen, USA) followed by UV cross-linking. The membrane was finally incubated with CSPD chemiluminescent solution and exposed to X-ray blue film (Retina, India).

### Salt stress treatment

Salt stress analysis was performed by using the young leaves from 2^nd^ and 3^rd^ nodes from 1-month-old *NtHY5OX*, *NtHy5CR* and WT plants to determine the comparative response during the leaf discs’ salt assay. These leaves were transferred into salt solution of 200 mM. The treatment was carried out at 25 °C for 3 days. In order to determine the effect of salinity stress, survival of leaves and chlorophyll content of NaCl treated leaves were analysed. All the data were recorded from 0 to 3 day of NaCl treatment. The experiment was repeated three times.

### Nitro blue tetrazolium (NBT) and 3,3′-diaminobenzidine (DAB) staining

Superoxide (O_2_-) staining was performed by infiltration by the nitro blue tetrazolium (NBT) method (Liu et al., 2018; Song et al., 2019) with some modification. WT as well as transgenic seedlings were grown for 10 days on half strength MS media then transferred to 200 mM salt solution for 3h. After 3 hours of salt stress seedlings were dipped in 0.5 mg/ml NBT solution for 2h followed by removal of chlorophyll by heating at 70°C temperature until chlorophyll is removed. Leaves were directly visualized for NBT staining.

Hydrogen peroxide (H_2_O_2_) staining of 10-day light grown tobacco seedlings was performed by staining with 3,3′-diaminobenzidine (DAB) using an adaptation of previous methods (Zhao et al 2016). WT as well as transgenic seedlings were grown for 10 days on half strength MS media then transferred to 200 mM salt solution for 3h. After 3 h of salt stress, seedlings were dipped in 1mg/ml DAB solution for 5 hours. Following the incubation, DAB staining solution was replaced with bleaching solution (ethanol: acetic acid: glycerol = 3:1:1). After that chlorophyll was removed by heating at 90-95°C temperature. Leaves were directly visualized for DAB staining.

### Statistical analysis

Data are plotted as means±SE (n=3) in the figures. Data of flavonol and nicotine accumulation were evaluated for statistical significance using the Two-tailed Student’s t-test using Graphpad prism 5.01 software. Asterisks indicate significance levels with (* P < 0.1; ** P < 0.01; *** P < 0.001) as confidence intervals.

## Author contributions

P.K.T. designed and supervised this study. D.S. designed and performed most of the experiments; HS, NS and SD provided help in conducting various experiments; P.K.T. analyzed the data; D.S., P.K.T. wrote the article; All authors read, contributed, and approved the article.

## Acknowledgment

P.K.T. acknowledges CSIR and DBT, New Delhi, for financial support in the form of projects on pathway engineering and genome editing. P.K.T. also acknowledges Department of Science and Technology, New Delhi for JC Bose National Fellowship. DS acknowledge University Grant Commission, New Delhi for Senior and Junior Research Fellowship respectively.

## Supplemental Data

**Supplemental Figure S1.** Estimation of flavonol content in WT tobacco.

**Supplemental Figure S2.** Sequence analysis of NtHY5 gene.

**Supplemental Figure S3.** Phylogenetic tree of NtHY5.

**Supplemental Figure S4.** Estimation of flavonol content in WT tobacco.

**Supplemental Figure S5.** Tissue-specific expression of NtHY5 and nicotine pathway genes.

**Supplemental Figure S6.** In vitro interaction between NtHY5 and the ACGT motif of G-BOX present at the promoters of *NtMYB12*.

**Supplemental Figure S7.** Complementation of *Arabidopsis hy5* mutant by NtHY5.

**Supplemental Figure S8.** Position of gRNA in NtHY5 sequences and amino acid sequence of truncated protein in mutants.

**Supplemental Figure S9.** NtHY5 modulates the gene expression of flavonoid biosynthesis genes.

**Supplemental Figure S10.** NtHY5 modulates flavonoid content in tobacco.

**Supplemental Figure S11.** NtHY5 leads to alteration in accumulation of anthocyanin content.

**Supplemental Figure S12.** Light responsiveness of NtHY5 and flavonol pathway genes.

**Supplemental Figure S13.** NtHY5 moves from shoot to root to regulate nicotine biosynthesis.

**Supplemental Figure S14.** NtHY5 regulates various morphological characterstics of tobacco seedlings.

**Supplemental Figure S15.** Comparative analysis of HY5OX vs. *NtHY5^CR^*.

**Supplemental Figure S16.** Effect of salt stress on NtHY5 transgenic lines of tobacco.

**Supplemental Figure S17.** Measurement of antioxidant activity in NtHY5OX and *NtHY5^CR^* lines in comparison to WT.

**Supplemental Table S1**. Putative cis-acting light responsive elements in NtPMT promoter.

**Supplemental Table S2.** Putative cis-acting light responsive elements in NtQPT promoter. **Supplemental Table S3**. Putative cis-acting light responsive elements in NtODC promoter. **Supplemental Table S4.** Putative cis-acting light responsive elements in NtMYB12 promoter.

**Supplemental Table S5.** Oligonucleotides used for development of constructs and expression analysis.

## Gene accession numbers

NtHY5 (XM_009590053), NtMYB12 (XM_016624824), NtODC (NW_017671182), NtQPT (NW_017672768), NtPMT (NW_017670241), NtTUB (XM_009622503), AtTUB (NM_125664).

## Data availability

All data generated or analyzed during this study are included in this published article (and its supplementary information files).

