## Supplementary Figures and Tables for "Mutation in shoot-to-root mobile transcription factor, ELONGATED HYPOCOTYL 5, leads to low nicotine levels in tobacco"

**Short title:** Light regulates nicotine biosynthesis

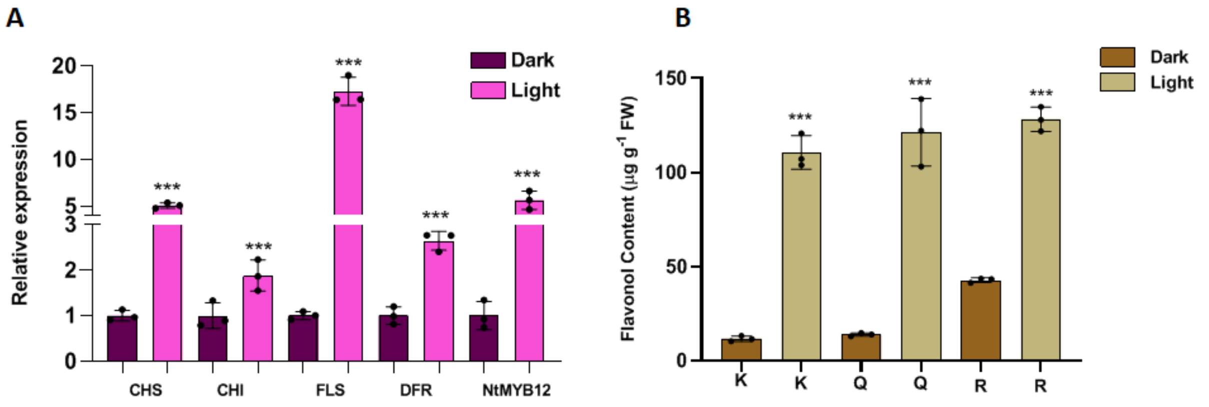


**Supplemental Figure S1. Estimation of flavonol content in WT tobacco.** A, Expression analysis of NtCHS, NtCHI, NtFLS, NtDFR and NtMYB12 in 10-d-old dark-grown and 10-d-old light-grown WT tobacco seedlings. B, Flavonol aglycones (K=kaempferol, Q =quercetin and R= rutin) were measured in 10-d-old dark-grown and 10-d-old light-grown WT tobacco seedlings. Tubulin was used as the endogenous control to normalize the relative expression levels. Statistical analysis was performed using two-tailed Student's t-test. Data are plotted as means ±SD (n=3). Error bars represent standard deviation. Asterisks indicate a significant difference, **P < 0.01, ***P < 0.001.

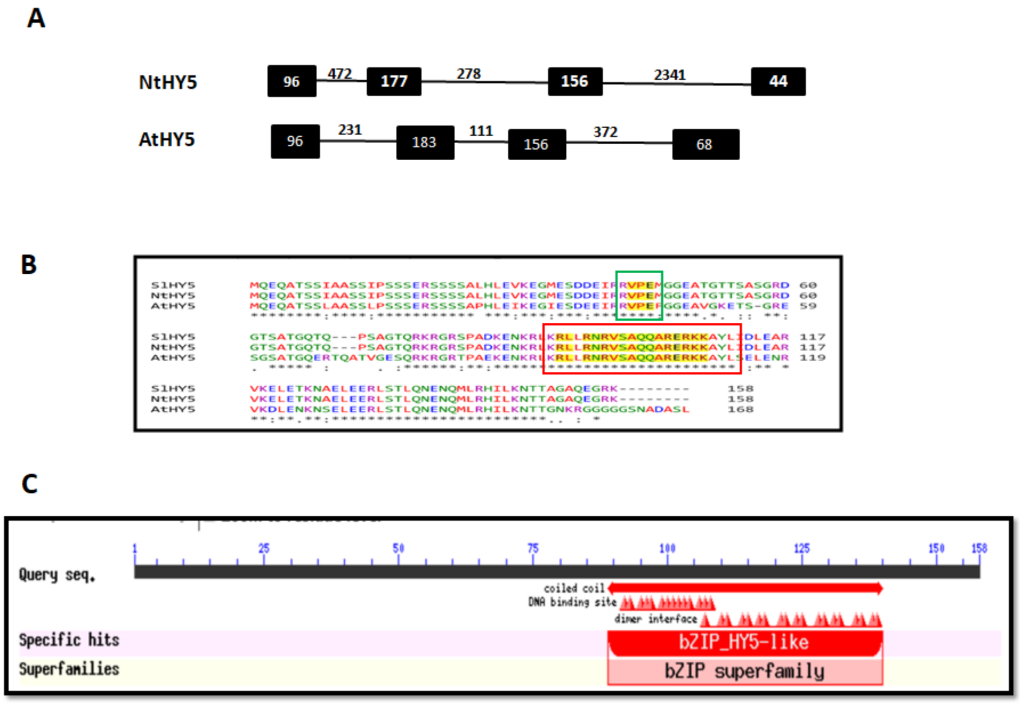


**Supplemental Figure S2. Sequence analysis of NtHY5 gene.** A, Gene structure of NtHY5 in comparison to AtHY5. The black boxes represent exons, lines represent introns and numbers represent the length of the exons and introns (bps). B, Amino acids alignment of NtHY5 and its homologs in Tomato and Arabidopsis using multallignment tool. The VPE motif and zinc finger domain is boxed in green and red color respectively. C, Representation of NtHY5 conserved domain that is basic leucine zipper (bZIP) domain through conserved domain database.

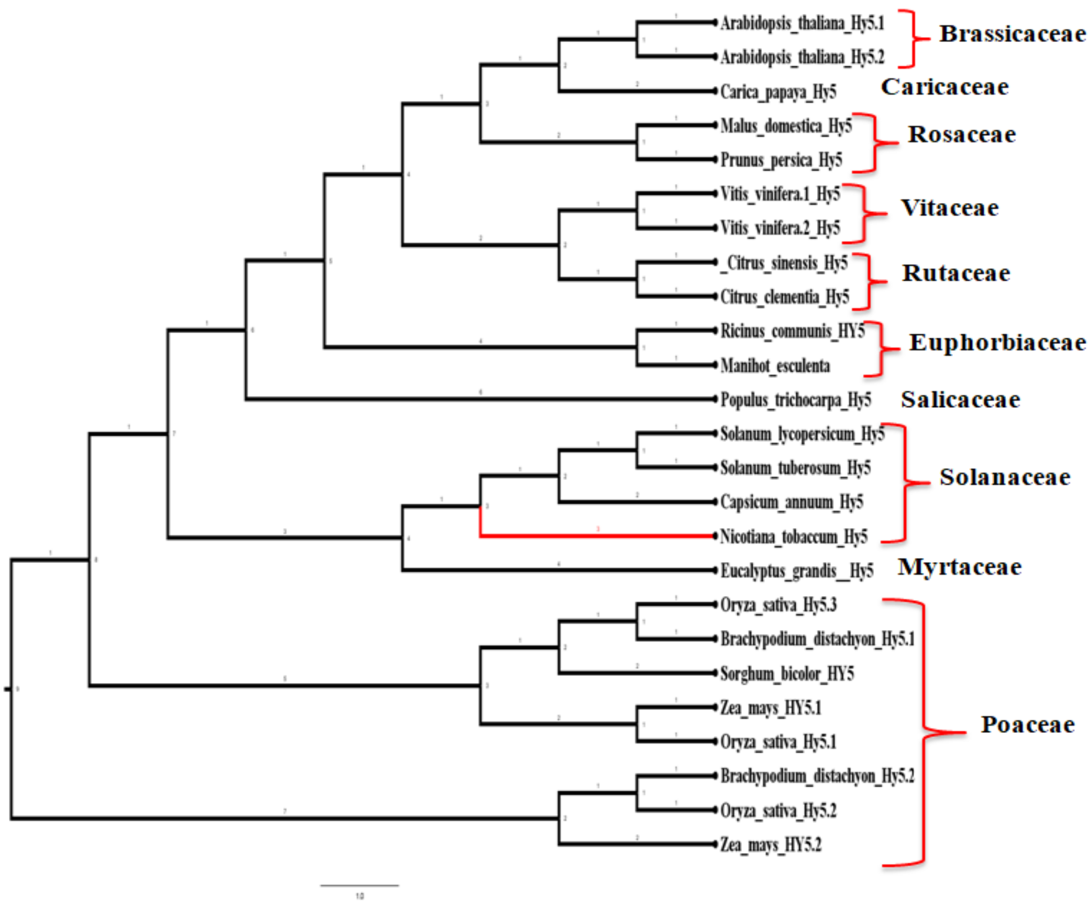


**Supplemental Figure S3. Phylogenetic tree of NtHY5.** Homologous proteins (sequence obtained from NCBI database) from across species of different families. The numbering at the nodes represents the number of trees that have the same topology as the consensus tree. NtHY5 is denoted through the red branch, and the scale bar indicates the branch length.

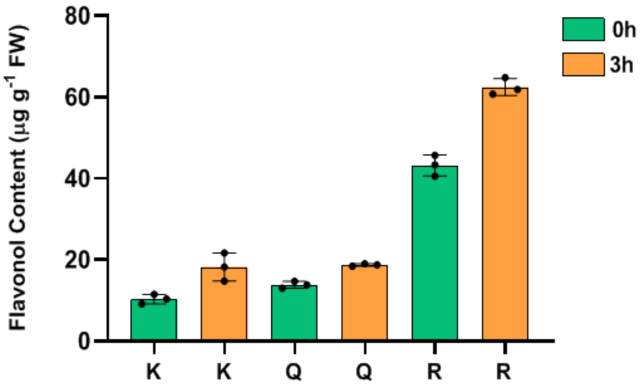


**Supplemental Figure S4. Estimation of flavonol content in WT tobacco.** Flavonol aglycones (K=kaempferol, Q=quercetin and R=rutin) were measured in 10-d-old dark-grown WT tobacco seedlings and WT seedlings transferred to light for 3 h after growing 10 days in darkness. Statistical analysis was performed using two-tailed Student's t-test. Data are plotted as means ±SD (n=3). Error bars represent standard deviation. Asterisks indicate a significant difference, **P < 0.01, ***P < 0.001.

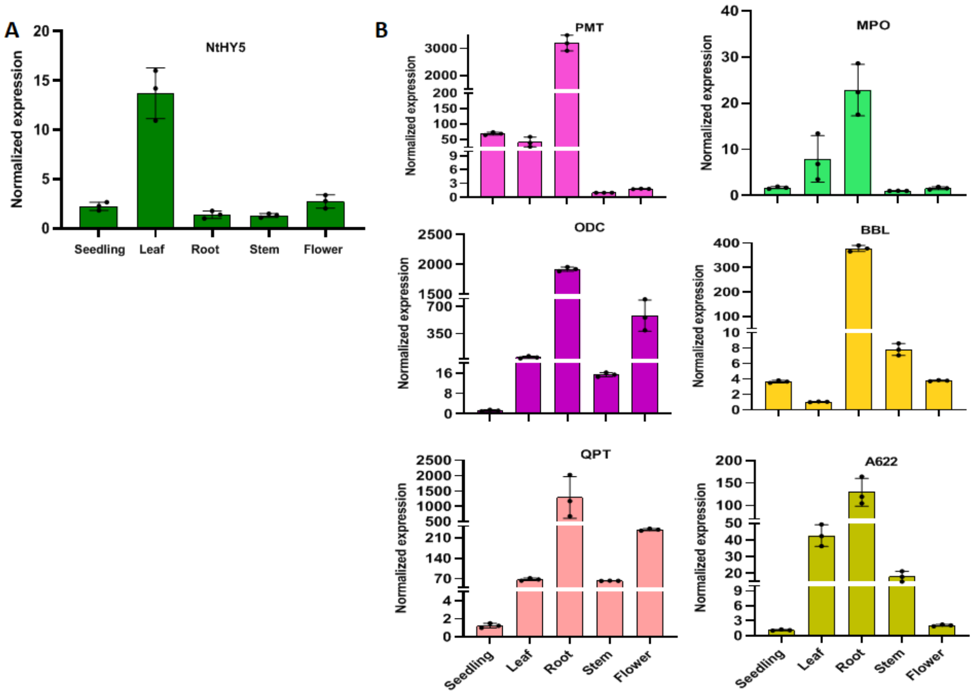


**Supplemental Figure S5**. **Tissue-specific expression of NtHY5 and nicotine pathway genes.** Normalized expression of A, NtHY5 and B-F, NtODC, NtPMT, NtMPO, NtA622 and NtQPT respectively in different tissue [seedling, leaf, stem, root and flower]. Tubulin was used as the endogenous control to normalize the relative expression levels. The statistical analysis was performed using two-tailed Student's t-tests. The data are plotted as means ± s.d (n=3). The error bars represent standard deviations.

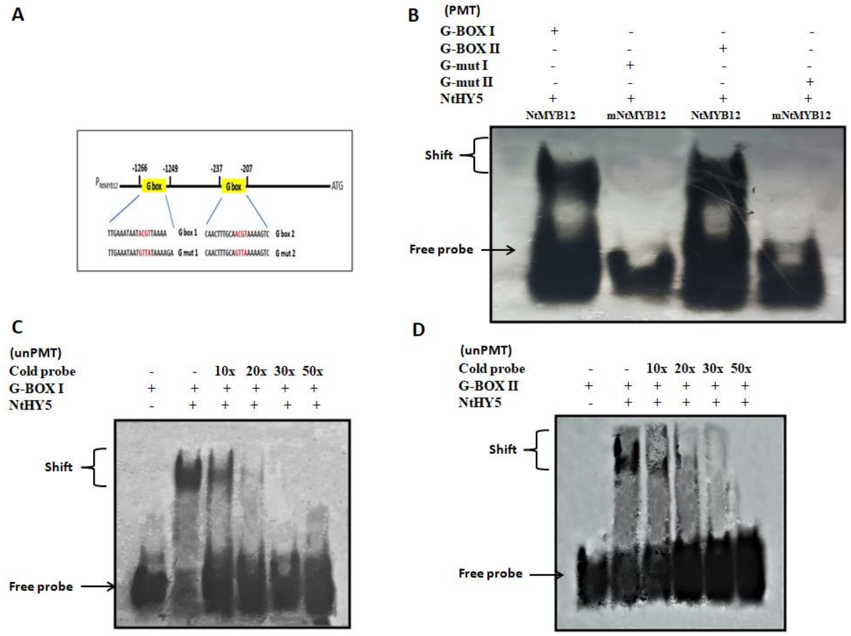


**Supplemental Figure S6. In vitro interaction between NtHY5 and the ACGT motif of G-BOX present at the promoters of *NtMYB12*.** A, Representation of the probes (G-BOXI, G-mut I, G-BOXII and G-mut II) designed upstream of the transcription start site of NtMYB12. G-BOX is altered by multiple base substitutions. B, EMSA (Electrophoretic mobility shift assay) for the binding of 6X-His-HY5 (NtHY5) with core ACE-DIG element (HY5 binding site that is G-BOXI and G-BOXII) present in the LRE motifs of the NtMYB12. Upper and lower arrows indicate shift and free probe respectively. C and D, EMSA with competition between digoxigen in labelled G-BOXI, unlabeled G-BOX I (cold), G-BOXII and unlabeled G-BOX II (cold) probe to bind to 6X-His-HY5 respectively. Superscript values 10X, 20X, 30X and 50X represent increasing amount of cold probe NtMYB12. Upper and lower arrows indicate shift and free probe respectively.

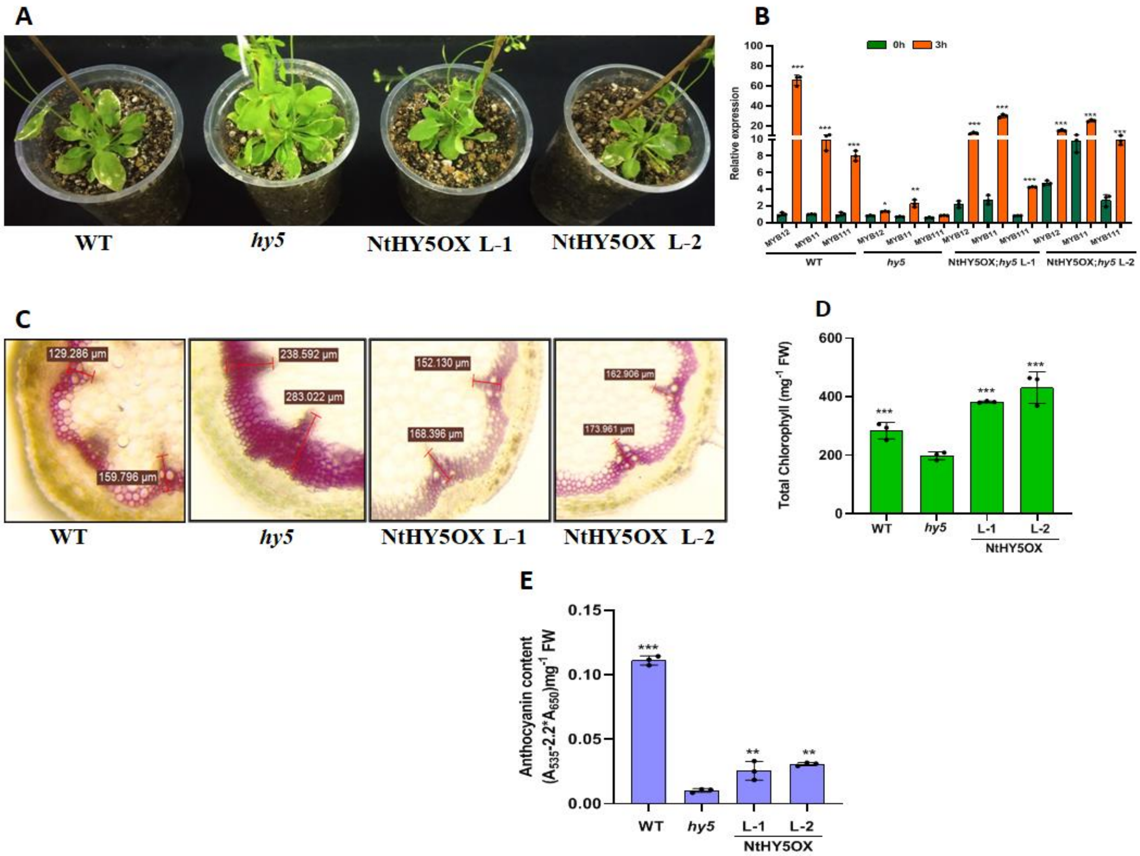


**Supplemental Figure S7. Complementation of Arabidopsis *hy5* mutant by NtHY5.** A, Representative photographs Rosette diameter of 30-day-old WT, *hy5*, L1 and L2 plants. **B,** Expression analysis of AtMYB12, AtMYB11 and AtMYB111 through real time PCR in NtHY5;*hy5*, hy5 and WT after growing for 7-d-dark condition and then transferred to light for 3 hours. C, Transverse section of stem to show lignin content in NtHY5; hy5 in comparison to hy5 and WT. D, Total Chlorophyll content in 30-d-old rosette leaves of WT, hy5 and NtHY5; hy5 plants. E, Anthocyanin content in WT, hy5 and NtHY5OX; hy5 in 7-d-old light-grown seedlings. Statistical analysis was performed using two-tailed Student's t-test. Tubulin was used as endogenous control to normalize the relative expression levels. Error bars represent ± s.d (n=3). For seed size (n=15). Asterisks indicate a significant difference, *P < 0.1, **P < 0.01, ***P < 0.001.

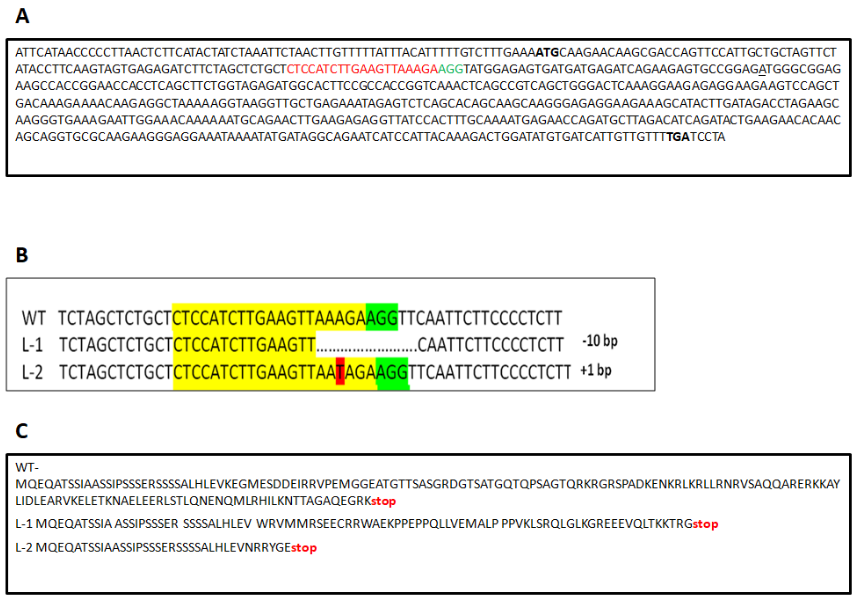


**Supplemental Figure S8. Position of gRNA in NtHY5 sequences and amino acid sequence of truncated protein in mutants.** A, Coding sequence of NtHY5 showing location of gRNA in red color, PAM sequences (NGG) are green in color. B, Schematic representation of nucleotide deletion/insertion in *NtHY5*$^{CR}$ edited tobacco plants. C, Amino acid sequence of WT and *NtHY5CR* edited plants.

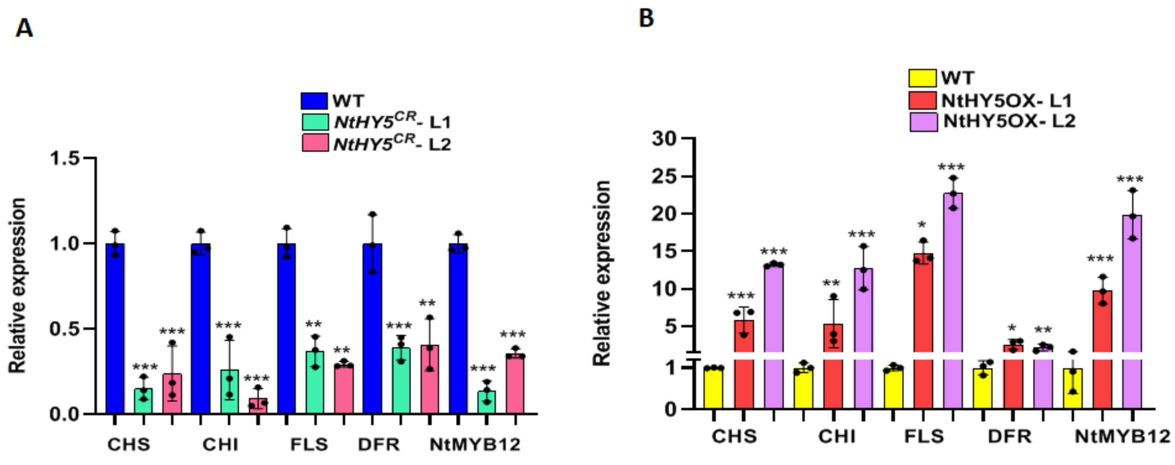


**Supplemental Figure S9. NtHY5 modulates the gene expression of flavonoid biosynthesis genes.** A, Relative expression of pheylpropanoid pathway genes (*NtCHS, NtCHI, NtFLS, NtDFR, and NtMYB12*) in 10-d-old light grown WT and *NtHY5*$^{CR}$ lines. B, Relative transcript abundance of pheylpropanoid pathway genes (*NtCHS, NtCHI, NtFLS, NtDFR and NtMYB12*) in 10-d-old light grown WT and NtHY5OX lines. Statistical analysis was performed using two-tailed Student's t-test. Error bars represent SE of means (n=3). Tubulin was used as endogenous control to normalize the relative expression levels. Asterisks indicate a significant difference, ***P < 0.001.

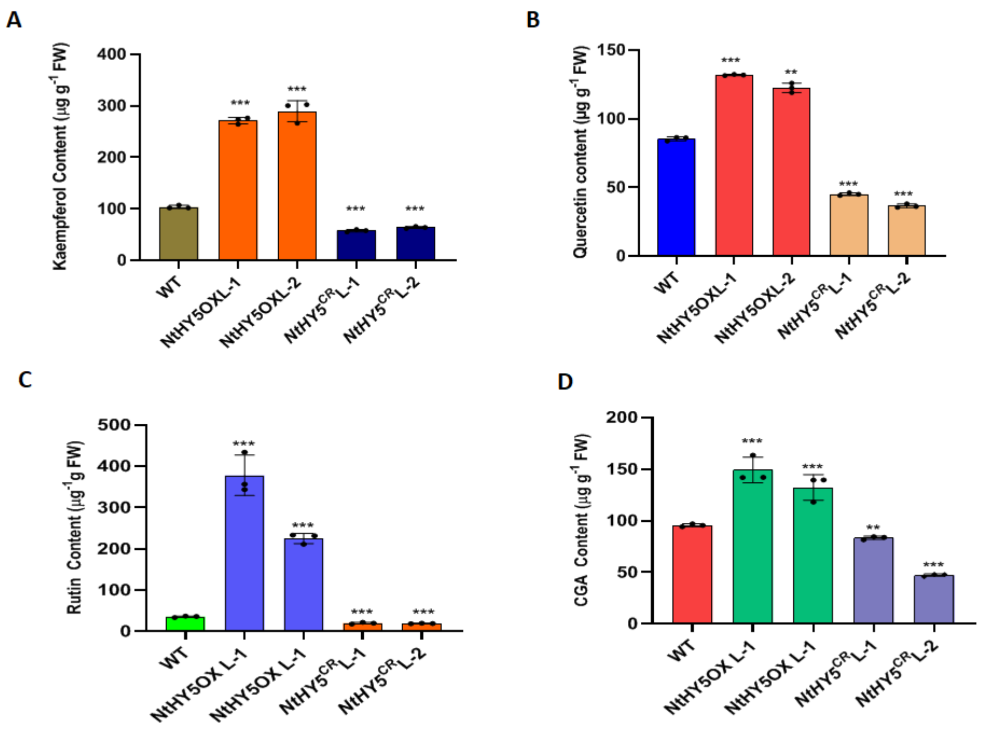


**Supplemental Figure S10. NtHY5 modulates flavonoid content in tobacco.** A-D, Quantification of flavonoids (K, Q, R) and polyphenolic compound (CGA) respectively in 10-d-old light grown WT, NtHY5OX and $\mathit{NtHY5}^{CR}$ lines of tobacco. Statistical analysis was performed using two-tailed Student's t-test. Error bars represent SE of means (n=3). Asterisks indicate a significant difference, ***P < 0.001.

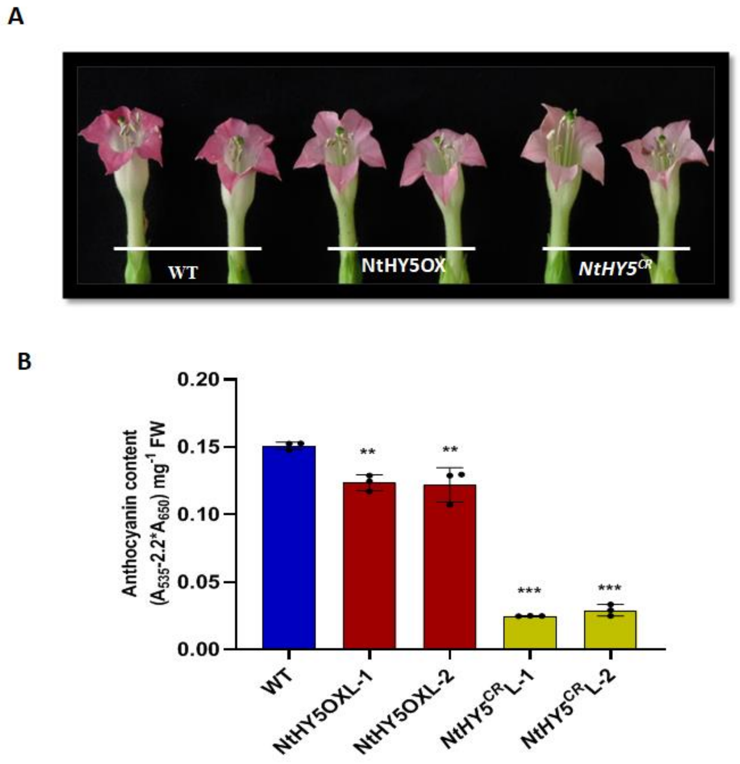


**Supplemental Figure S11. NtHY5 leads to alteration in accumulation of anthocyanin content.** A, Representative photograph of flower color alterations in petals of NtHY5OX and $NtHY5^{CR}$ transgenic lines in compared with WT. B and C, Quantification of Anthocyanin content in 10-d-old light grown WT, NtHY5OX and $NtHY5^{CR}$ lines. Statistical analysis was performed using two-tailed Student's t-test. Error bars represent SE of means (n=3). Asterisks indicate a significant difference, ***P < 0.001.

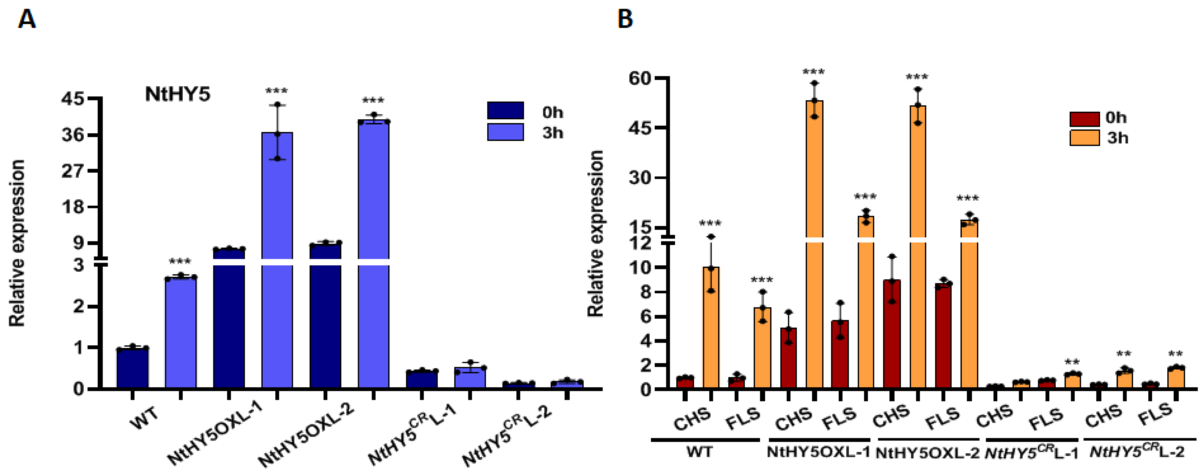


**Supplemental Figure S12. Light responsiveness of NtHY5 and flavonol pathway genes.** A, B, Relative expression of NtHY5 and phenylpropanoid pathway genes (*NtCHS*, *NtCHI*, *NtFLS* and *NtDFR*) respectively in 10-d-old dark grown seedlings transferred to light for 3h. Statistical analysis was performed using two-tailed Student's t-test. Error bars represent SE of means (n=3). Tubulin was used as endogenous control to normalize the relative expression levels. Asterisks indicate a significant difference, *P < 0.1, **P < 0.01, ***P < 0.001

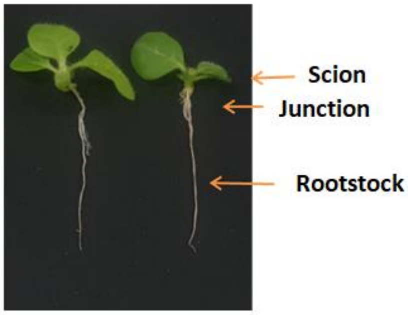


**Supplemental Figure S13.** Phenotype of grafted seedlings.

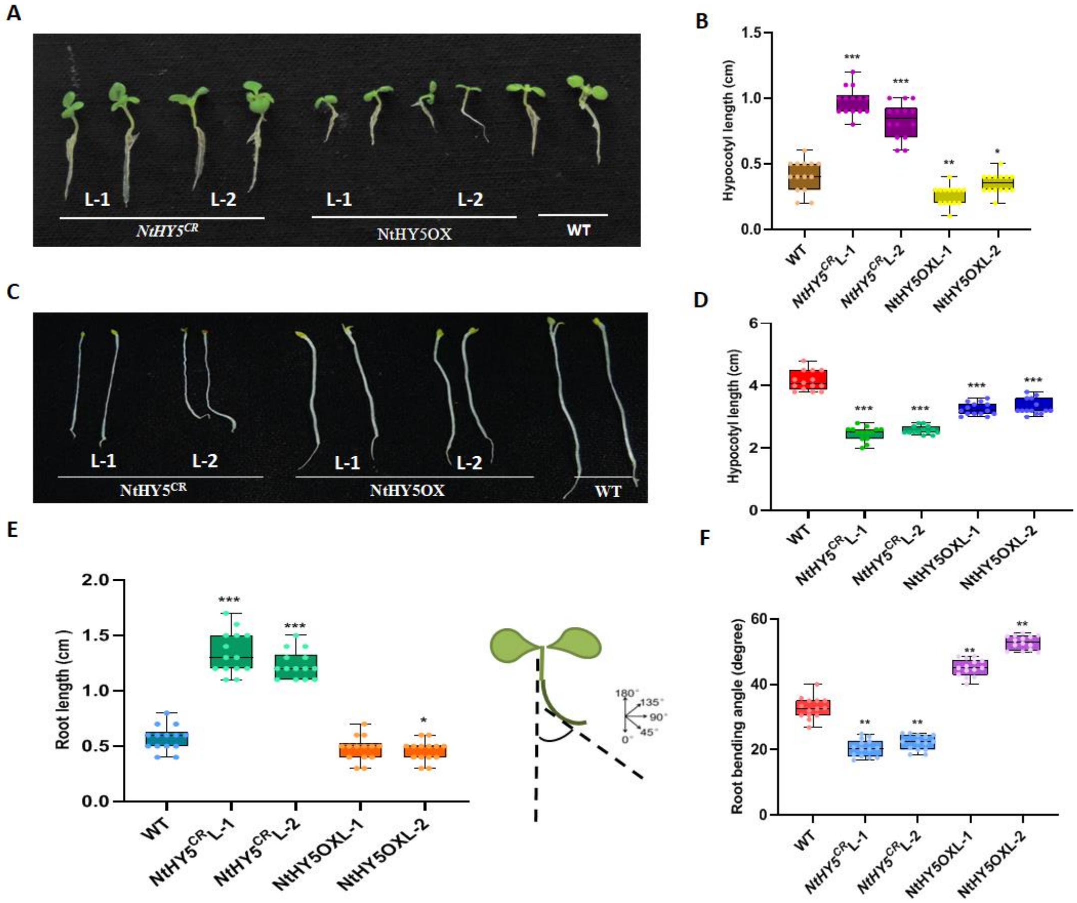


**Supplemental Figure S14. NtHY5 regulates various morphological characterstics of tobacco seedlings.** A, Phenotype of WT, *NtHY5*$^{CR}$ and NtHY5OX 10-d-light grown seedling. B, Measurement of hypocotyls length of WT, *NtHY5*$^{CR}$ and NtHY5OX 10-d-light grown seedling. C, Phenotype of WT, *NtHY5*$^{CR}$ and NtHY5OX 10-d-dark grown seedling. D, Measurement of hypocotyls length of WT, *NtHY5*$^{CR}$ and NtHY5OX 10-d-dark grown seedling. E, Root length of 10-d-old light grown WT, NtHY5OX and *NtHY5*$^{CR}$ seedlings. F, Measurement of root bending angle of 10-d-old light grown WT, NtHY5OX and *NtHY5*$^{CR}$ seedlings. Tubulin was used as the endogenous control to normalize the relative expression levels. The statistical analysis was performed using two tailed Student's t-tests. The data are plotted as means ± SD (n= 3). For measurement of root bending angle, root length, hypocotyls length (n=10-12). The error bar represents standard deviations. The asterisks indicate significant difference, *P< 0.1; **P< 0.01; 0.001.

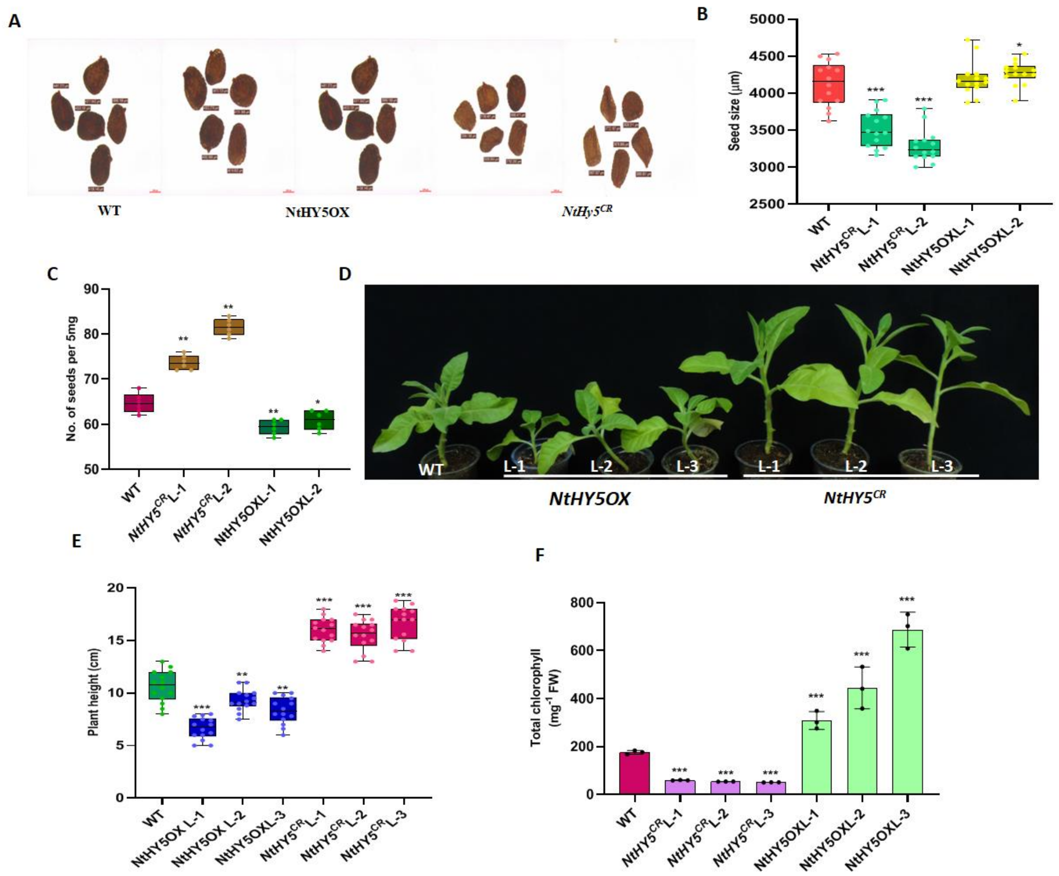


**Supplemental Figure S15. Comparative analysis of HY5OX vs. *NtHY5*$^{CR}$.** A, Representative image of seed size of NtHY5OX and *NtHY5*$^{CR}$ plants in comparison to WT. B, Measurement of seed size of NtHY5OX and *NtHY5*$^{CR}$ plants in comparison to WT. C, Number of seeds per 5mg of NtHY5OX and *NtHY5*$^{CR}$ plants in comparison to WT. D, Representative image of 40-day-old WT, NtHY5OX and *NtHY5*$^{CR}$ plants showing plant heights. E, Measurement of plant height of 40-day-old NtHY5OX and *NtHY5*$^{CR}$ young plants in comparison to WT. F, Quantification of total chlorophyll content of WT, NtHY5OX and *NtHY5*$^{CR}$ 10-day light grown seedlings. Statistical analysis was performed using two-tailed Student's t-test. Error bars represent SE of means (n=3). For seed size and hypocotyls length (n=15). Asterisks indicate a significant difference, $*P < 0.1$, $**P < 0.01$, $***P < 0.001$. Statistical analysis was performed using two-tailed Student's t-test. Data are plotted as means ±SD (n=3). Error bars represent standard deviation. Asterisks indicate a significant difference, $***P < 0.001$.

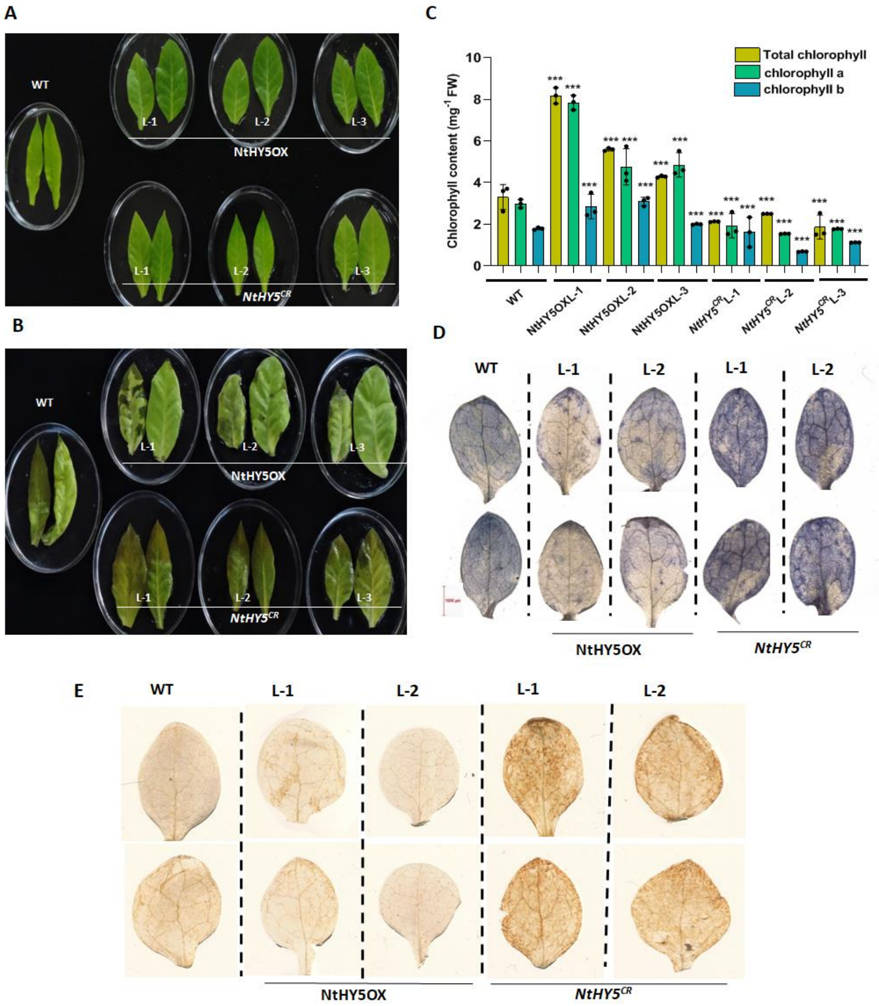


**Supplemental Figure S16. Effect of salt stress on NtHY5 transgenic lines of tobacco.** Fully expanded leaves of 30-d-old plants to floated on 6 ml of 200 mM NaCl solution (for salt stress) for 3 days. A and B, The phenotypic (chlorosis) changes due to salt treatment on leaf discs were assessed on 0 and 3 days respectively. C, Quantification of chlorophyll content of WT, NtHY5OX and *NtHY5$^{CR}$* leaves after 3day of salt stress. D, Detection of Superoxide ($O^{2-}$) by NBT staining in 10-d-old light grown WT, NtHY5OX and *NtHY5$^{CR}$* tobacco seedlings subjected to 200 mM salt stress for 3 hours. E, Detection of hydrogen peroxide (H2O2) by DAB staining in 10-d-old light grown WT, NtHY5OX and *NtHY5CR* tobacco seedlings. The statistical analysis was performed using two-tailed Student's t-tests. The data are plotted as means ± SD (n=3). The error bars represent standard deviations. The asterisks indicate significant difference, *P< 0.1; **P< 0.01; ***P< 0.001.

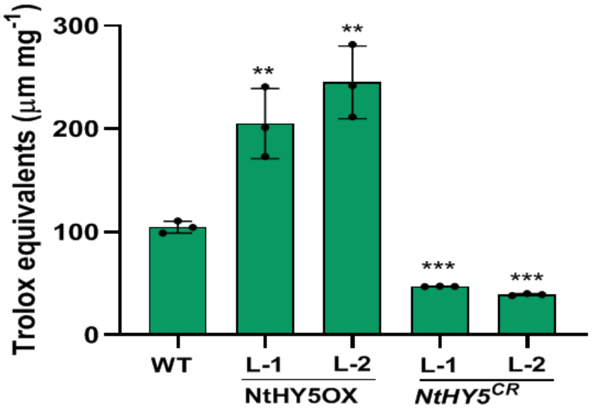


**Supplemental Figure S17. Measurement of antioxidant activity in** NtHY5OX and ***NtHY5**$^{CR}$* **lines** in comparison to WT**.** Estimation of antioxidant activity by trolox equivalent in WT, NtHY5OX and *NtHY5*$^{CR}$ lines. Statistical analysis was performed using two-tailed Student's t-test. Error bars represent SE of means (n=3). Asterisks indicate a significant difference, **P < 0.01, ***P < 0.001.

**Supplemental Table S1**. Putative cis-acting light responsive elements in NtPMT promoter.

| Site Name | Sequence | Position | Function |
|---|---|---|---|
| ACE | CTAACGTATT | -1967 | cis-acting element involved in light responsiveness |
| G-box | CACGTT | -184<br>-1057<br>-1732 | cis-acting regulatory element involved in light responsiveness |
| I-box | TGATAATGT | -1069 | part of a light responsive element |
| TCCC motif | TCTCCCT | -389 | part of a light responsive element |
| TCT motif | TCTTAC | -250<br>-1923 | part of a light responsive element |

**Supplemental Table S2.** Putative cis-acting light responsive elements in NtQPT promoter.

| Site Name | Sequence | Position | Function |
|---|---|---|---|
| AE-box | AGAAACAA | -705 | cis-acting element involved in light responsiveness |
| Box-4 | ATTAAT | -517<br>-897<br>-1580 | part of a conserved DNA module involved in light responsiveness |
| G-box | CACGTT | -56 | cis-acting regulatory element involved in light responsiveness |
| GA motif | ATAGATAA | -1722 | part of a light responsive element |
| GATA motif | GATAGGA | -712 | part of a light responsive element |
| GT1 motif | GGTTAAT | -1994 | light responsive element |
| LAMP element | CTTTATCA | -1821 | part of a light responsive element |

**Supplemental Table S3**. Putative cis-acting light responsive elements in NtODC promoter.

| Site Name | Sequence | Position | Function |
|---|---|---|---|
| Box-4 | ATTAAT | -302<br>-1678<br>-1752<br>-1811<br>-1986 | part of a conserved DNA module involved in light responsiveness |
| G-box | CACGTT | -194 | cis-acting regulatory element involved in light responsiveness |
| GT1 motif | GGTTAAT | -1737 | light responsive element |
| Gap-box | CAAATGAA(A/G)A | -1273 | part of a light responsive element |

**Supplemental Table S4.** Putative cis-acting light responsive elements in NtMYB12

| Site Name | Sequence | Position | Function |
|---|---|---|---|
| Box-4 | ATTAAT | -503<br>-1481<br>-1561 | part of a conserved DNA module involved in light responsiveness |
| ACE | CTAACGTATT | -1255 | cis-acting element involved in light responsiveness |
| GATA motif | AAGATAAGATT | -1245 | light responsive element |

**Supplemental Table S5.** Oligonucleotides used for development of constructs and expression analysis.

| S.No. Gene | | Forward Primer (5' to 3') | Reverse Primer(5' to 3') |
|---|---|---|---|
| 1. | **NtHY5** | TTTTG**GGATCC**AAAATGCAAGAACAAGC | GAGGAAATAAAAT**GAGCTC**GGCAGAAT |
| 2. | **PETNtHY5** | TTTGTC**GGATCC**AATGCAAGAACAA | GAGGAAATAAAAT**GAGCTC**GGCAGAAT |
| 3. | **NtHY5gRNA** | ATTGCTCCATCTTGAAGTTAAAGA | AAACTCTTTAACTTCAAGATGGAG |
| 4. | **NtQPT P1** | CATAAATAA**AACGTG**TTCAGCTACTAAAAC | GTTTTAGTAGCTGAACACGTTTTATTTATG |
| 5. | **NtQPT Mut P1** | CATAAATAA**AATTCA**TTCAGCTACTAAAAC | GTTTTAGTAGCTGAATGAATTTTATTTATG |
| 6. | **NtODC P1** | ATCTGATCACCT**AACGTG**ACCAAGAAAATA | TATTTTCTTGGTCACGTTAGGTGATCAGAT |
| 7. | **NtODC Mut P1** | ATCTGATCACC**TAATATC**ACCAAGAAAATA | TATTTTCTTGGTGATATTAGGTGATCAGAT |
| 8. | **NtPMT P1** | CAGTCTAACCATG**CACGTT**GTAATGATTTT | AAAATCATTACAACGTGCATGGTTAGACTG |
| 9. | **NtPMTP2** | TACTGTTTCCGC**AACGTG**CTCTTCATCAGG | CCTGATGAAGAGCACGTTGCGGAAACAGTA |
| 10. | **NtPMT Mut P1** | CAGTCTAACCATG**ATTAAT**GTAATGATTTT | AAAATCATTACATTAATCATGGTTAGACTG |
| 11. | **NtPMT Mut P2** | TACTGTTTCCGC**AATCTT**CTCTTCATCAGG | CCTGATGAAGAGAAGATTGCGGAAACAGTA |
| 12. | **NtMYB12 P1** | AACTTGAAATAAT**ACGT**TAAAAGATAAAA | TTTTATCTTTTAACGTATTATTTCAAGT |
| 13. | **NtMYB12 Mut P1** | AACTTGAAATAAT**GTTA**TAAAAGATAAAAT | ATTTTATCTTTTATAACATTATTTCAAGTT |
| 14. | **NtMYB12 P2** | AGTCAACTTTGCA**ACGT**AAAAGTCCACACA | TGTGTGGACTTTTACGTTGCAAAGTTGACT |
| 15. | **NtMYB12 Mut P2** | AGTCAACTTTGCA**GTTA**AAAAGTCCACACA | TGTGTGGACTTTTTAACTGCAAAGTTGACT |
| 16. | **M13** | GTAAAACGACGGCCAGT | CAGGAAACAGCTATGAC |
| 17. | **CaMV35S** | GTAAGGGATGACGCACAATCC | GGACTCTAATCATAAAAACCC |
| 18. | **Tubulin** | GAGCCTTACAACGCTACTCTGTCTGTC | ACACCAGACATAGTAGCAGAAATCAAG |
| 19. | **pHSE401** | TGTCCCAGGATTAGAATGATTAGGC | CCAGAAATTGAACGCCGAAGAAC |
| 20. | **RTNtPMT** | CCGAACAACAGAACGGGACAA | GAGAATGCTTCACCTGGCCAT |
| 21. | **RT NtQPT** | GCAAAGAATGAGTGGAATAGCT | ATCAATACCGCCCATTTATCC |
| 22. | **RT NtODC** | CATCACCCGAAATCCGAACTC | GGCTCGACTTCTTCTGGAAGC |
| 23. | **RT NtMPO** | GTGAAGCTGTGGTGAAAAGTTA | GGAGCATCAGCCTCACTATG |
| 24. | **RT NtA622** | GGATAGAATATTGGCAGG | GGATAGAATATTGGCAGGTGG |
| 25. | **RT NtCHS** | AACTAGACTCCAATTTCTGGAATG | AGCCCAGGAACATCTTTGAG |
| 26. | **RT NtCHI** | GTCAGGCCATTGAAAAGCTC | CTAATCGTCAATGCCCCAAC |
| 27. | **RT NtFLS** | TTTGGCACTTGGTGTTGTGG | ACTTGACATCATACCAATGGC |

| | | |
|---|---|---|
| **28. RT NtMYB12** | CTGAAAACCGACCAATCCGT | CCGATCGAAGTGGGTTGGTA |
| **29. RT AtMYB111** | AACCAACAGTCAGGGGAATG | TTGGACATCGACAGTTCCAG |
| **30. RT AtMYB12** | TCAGCCGTAAACTCCACAACTTC | CTCAAGACGTCTCCGCCG |
| **31. RT AtMYB11** | CAGTCCAACGGCGAAGGA | CAAAAATGCCGGGCTAAAGA |
| **32. RT AtCHS** | AAGCGCATGTGCGACAAG | CACATGCATCTGACGGAGGA |
| **33. RT AtCHI** | CCGGTTCATCGATCCTCTTC | CTTACGGTTGCGTTTTCGAAA |
| **34. RT AtFLS** | TCACAACATTCCGAGGTCCAA | TCGATCTAAGCGATCCCGAC |
| **35. RT AtDFR** | TGGTGGTCGGTCCATTCAT | CCTTATCACCGCGCTCTCTC |

Bold with under line sequences are restriction sites incorporated in the sequences for cloning, bold sequences are G-BOX core motif and mutated motif used in EMSA.
